# Endogenous modulation of pain relief: evidence for dopaminergic but not opioidergic involvement

**DOI:** 10.1101/2022.07.10.499477

**Authors:** Simon Desch, Petra Schweinhardt, Ben Seymour, Herta Flor, Susanne Becker

## Abstract

Relief of ongoing pain is a potent motivator of behavior, directing actions to escape from or reduce potentially harmful stimuli. Whereas endogenous modulation of pain events is well characterized, relatively little is known about the modulation of pain relief and its corresponding neurochemical basis. Here we studied pain modulation during a probabilistic relief-seeking task (a ‘wheel of fortune’ gambling task), in which people actively or passively received reduction of a tonic thermal pain stimulus. We found that relief perception was enhanced by active decisions and unpredictability, and greater in high novelty-seeking trait individuals, consistent with a model in which relief is tuned by its informational content. We then probed the roles of dopaminergic and opioidergic signaling, both of which are implicated in relief processing, by embedding the task in a double-blinded cross-over design with administration of the dopamine precursor levodopa and the opioid receptor antagonist naltrexone. We found that levodopa, but not naltrexone, enhanced each of these information-specific aspects of relief modulation. These results show that dopaminergic signaling has a key role in modulating the perception of pain relief to optimize motivation and behavior.

## Introduction

When we are in pain, our desire for pain relief and the pleasure of pain relief are universally appreciated. However, research into the state of pain has gained considerably more attention than that of relief. Theoretical perspectives on pain typically focus on its aversiveness, reflecting the powerful incentive to avoid harm wherever possible. Perceived pain is highly sensitive to the motivational context, with modulatory processes appearing to endogenously tune pain perception to help optimize the way in which it controls responses and action (Fields, 2018; Seymour, 2019). However if pain is ongoing, there is an equally potent new incentive to reduce or escape from it, in which pain relief arises as a strong positive motivational force and a reinforcement signal in its own right (Leknes, Brooks, Wiech, & Tracey, 2008; Becker, Gandhi, Kwan, Ahmed, & Schweinhardt, 2015). However, how relief acts as a signal for shaping behavior is less studied: in particular, it is not clear the extent to which the perception of relief is also sensitive to endogenous modulation; and if so, how such modulation is neurally mediated.

In general, endogenous modulation of pain involves a number of different processes mediated by distinct descending signaling pathways (Bannister, 2019). Opioid-based mechanisms are important for many of these. Work in rodents has shown that endogenous opioid activity in the anterior cingulate cortex is necessary and sufficient to induce the rewarding effects of relief from pain (Navratilova et al., 2015). In humans, the perceived pleasantness and magnitude of pain relief has been shown to decrease with administration of the opioid antagonist naltrexone, confirming a role of opioids in pain relief perception (Sirucek et al., 2021). Other forms of endogenous modulation, such as the placebo effect, are also opioid-sensitive (Benedetti, 1996; Eippert et al., 2009; King et al., 2013). Alongside this, however, dopaminergic-based mechanisms also have a clear role. For instance conditioned place preference induced by pain relief is associated with activity in midbrain dopaminergic neurons (Navratilova et al., 2015, 2012; Xie et al., 2014). In the case of primary rewards, dopamine is implicated in the active motivation to obtain reward (“wanting”), while endogenous opioids mediate the hedonic experience of reward (“liking”). (Barbano & Cador, 2006, 2007; Berridge, Robinson, & Aldridge, 2009; Sherdell, Waugh, & Gotlib, 2012; Smith, Berridge, & Aldridge, 2011; Tindell, Berridge, Zhang, Peciña, & Aldridge, 2005). However, the extent to which this distinction might hold for pain relief is not clear.

In theoretical models of pain motivation, endogenous modulation of pain is considered an action in its own right, with the pain system making active ‘decisions’ to tune incoming pain signals so as to optimize responding in a given situation (the ‘*Motivation Decision Model of Pain*’: Fields, 2006, 2007, 2018). One example of this is in inhibition of pain of external rewards, which allows suppression of immediate nocifensive responses that could interfere with more important goals. Studies in humans indicate that this is dopamine sensitive (Becker, Gandhi, Elfassy, & Schweinhardt, 2013). But whether purely endogenous modulation of pain relief is dopamine sensitive is not known, not least because relief modulation is not well characterized to begin with. Evidence does exist that active pain relief-seeking, when compared to passive relief receipt, is associated with enhanced pain relief perception, and this phenomenon is associated with novelty-seeking traits (Becker et al., 2015). This would fit with information-processing accounts of endogenous modulation (Seymour, 2019), which propose that pain is modulated to optimize prospective control of behavior. Whether this is sensitive to opioidergic or dopaminergic (or both) signaling is not known.

The aim of the present study was therefore first to better characterize information processing aspects of relief motivation, and second to investigate the roles of dopaminergic and opioidergic signaling. We expected that pain relief would be modulated by the value of information it carries, as hence enhanced by i) active vs passive reception and thus controllability, since this reflects potential to exploit relief information; ii) unpredictability, since this reflects the extra information carried by surprising events, and iii) trait novelty-seeking, since this reflects individual information sensitivity. At the same time, we aimed to identify the potential role of dopamine and opioids for each of these factors, in particular to explore whether increased dopamine availability would enhance endogenous pain relief under these conditions, and whether modulation could be reduced by blocking opioid receptors. Finally, we aimed to identify whether modulation of relief was also apparent in the explicit decisions that arise in probabilistic learning, to determine whether perception of relief can be dissociated from instrumental choice.

To test these hypotheses, we employed a previously developed wheel of fortune task utilizing relief of a tonic capsaicin-sensitive thermal pain stimulus as ‘wins’, and allowing to quantify endogenous pain inhibition induced by gaining pain relief in active versus passive conditions (Becker et al., 2015). To test the roles of dopamine and opioids, healthy volunteers ingested either a single dose of the dopamine precursor levodopa (150mg), the opioid antagonist naltrexone (50mg), or placebo in separate testing sessions (double-blinded, placebo controlled cross-over design). To allow also the assessment of reinforcement learning, a probabilistic reward schedule associated with the participants’ choices in the wheel of fortune was implemented.

## Results

### Endogenous modulation of active pain relief seeking under placebo

To test whether playing the wheel of fortune induced endogenous pain inhibition by gaining pain relief during active (controllable) decision-making, a test condition in which participants ‘won’ relief of a tonic thermal pain stimulus in the game was compared to a control condition with passive receipt of the same outcomes (Figure 1). As a further comparator the game included an opposite condition in which participants received *increases* of the thermal stimulation as punishment. This loss condition was also complemented by a passive condition involving receipt of the same nociceptive input.

**Figure 1:**
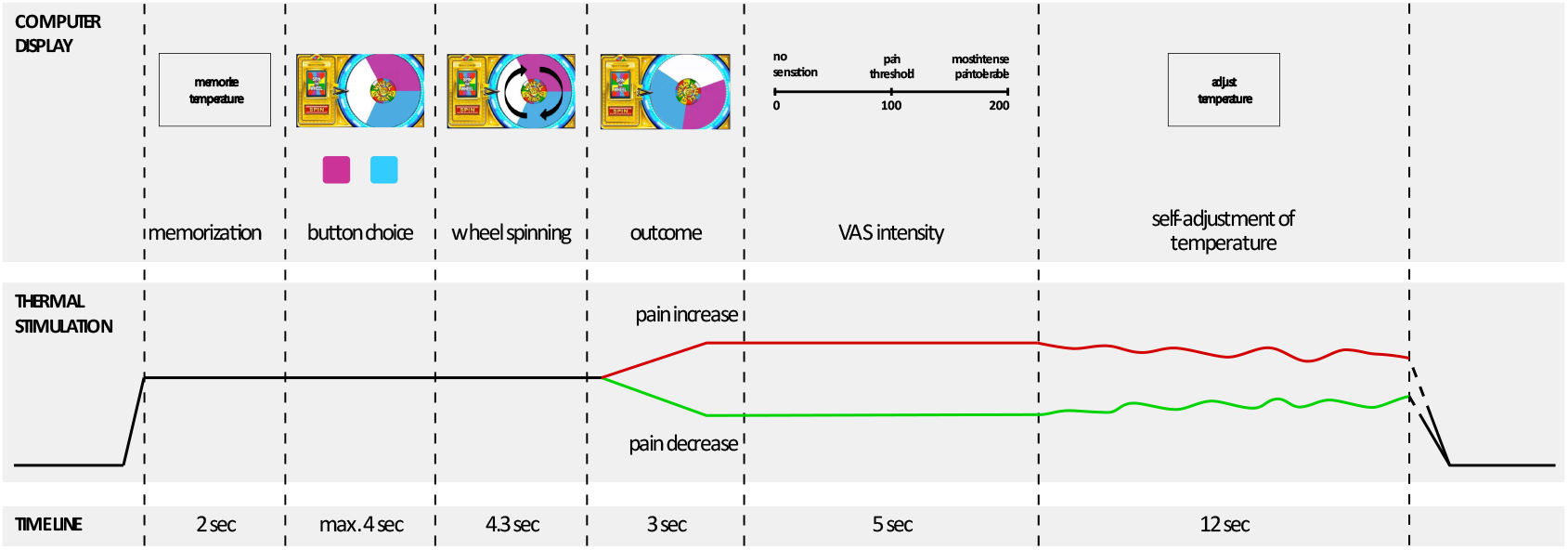
Time line of one trial with active decision-making (test trials) of the wheel of fortune game. In each test session, one of the two colors (pink and blue) of the wheel was associated with a higher chance to win pain relief (counterbalanced across subjects and drug conditions). Pain relief (win) as outcome of the wheel of fortune game is depicted in green, pain increase (loss) in red. In passive control trials and neutral trials subjects did not play the game but had to press a black button after which the wheel started spinning and landed on a random position with no pointer on the wheel. Trials with active decision-making were complemented by passive control trials without decision making but the same nociceptive input (control trials), resulting in the same number of pain increase and pain decrease trials as in the active condition. In neutral trials the temperature did not change during the outcome interval of the wheel. In all trial types, participants had to adjust the temperature to the memorized sensation at the beginning of the trial as an operationalization of a behavioral assessment of pain sensitization and habituation across the course of one trial. Adapted from (Becker et al., 2015).

#### Ratings of perceived pain

Replicating previous results, in the placebo (i.e. non-drug) condition participants rated the thermal stimulation as less intense after actively winning pain relief compared to the passive control condition, as rated on visual analogue scales (VAS) from “no sensation” (0) over “just painful” (100) to “most intense pain tolerable” (200). Furthermore, participants also rated the stimulation as more intense after actively losing compared to the passive control condition (Figure 2 A; interaction ‘outcome × trial type’, *F*(1,1040) = 64.14, *p* < 0.001; pairwise comparisons: win: test vs. control *p* < 0.001; lose: test vs. control, *p* < 0.001). This shows that perception of both relief and pain are enhanced by active (instrumental) controllability, as hypothesized.

**Figure 2:**
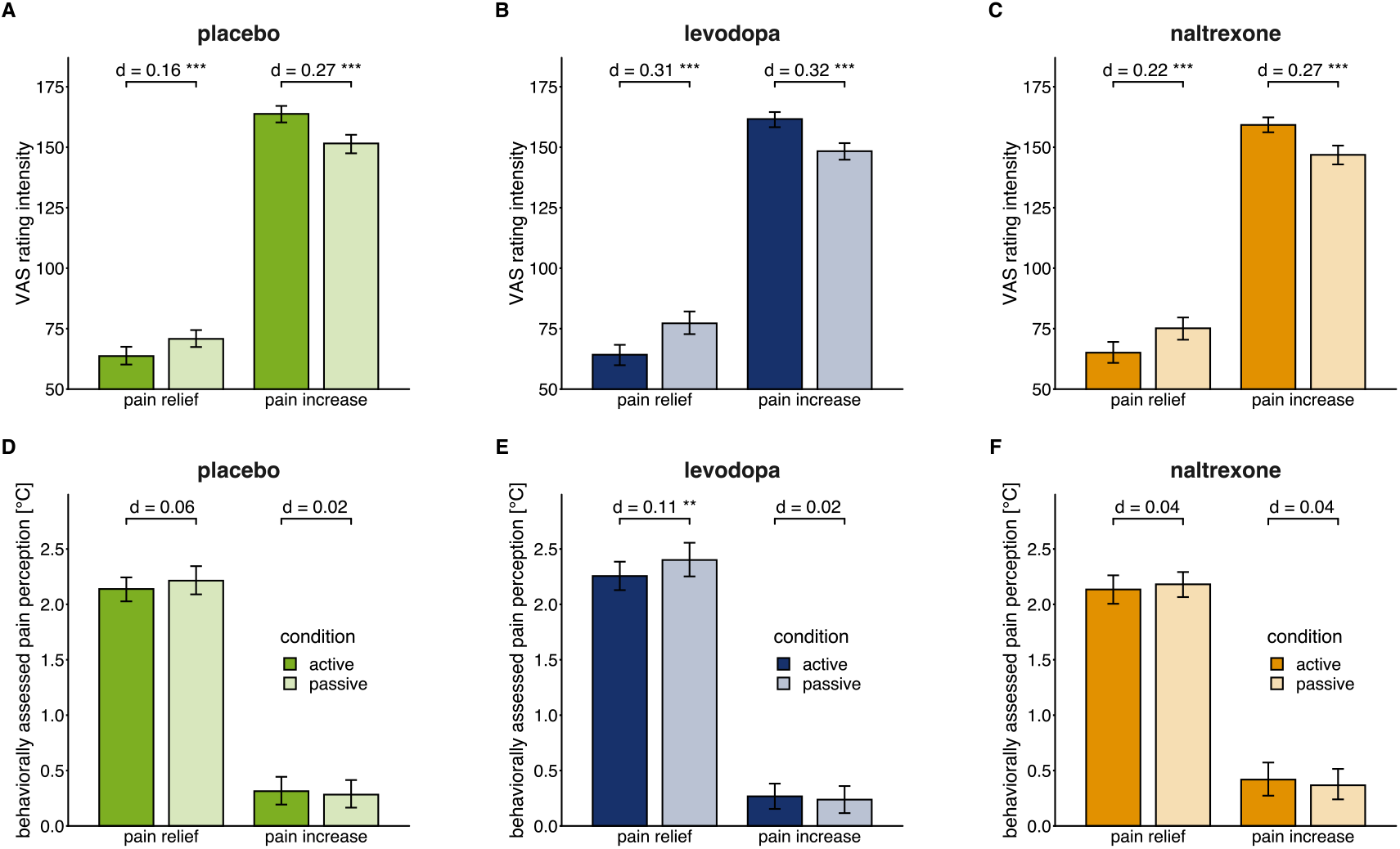
Means and 95% confidence intervals of means for VAS pain intensity ratings (A, B, C) and behaviorally assessed pain perception (D, E, F; within-trial sensitization in pain perception in °C) for each drug session (placebo: n = 28, levodopa: n = 27, naltrexone: n = 28). d indicates Cohen’s d as standardized effect-size of estimated effects. ** *p* < 0.01, *** *p* < 0.001, for post-hoc comparisons of test versus control trials.

#### Behaviorally assessed pain perception

In addition to the VAS ratings, participants performed a validated perceptual task (Becker, Kleinböhl, Baus, & Hölzl, 2011; Kleinböhl et al., 1999) allowing to assess perception of the underlying tonic pain stimulus, which is specifically sensitive to perceptual sensitization and habituation. In this procedure, participants re-adjust the stimulation temperature themselves after the outcome of the wheel of fortune to match their perception at the beginning of trial. Negative values (i.e. higher re-adjusted temperatures compared to the stimulation intensity at the beginning of the trial) indicate habituation across the course of one trial of the game, positive values indicate sensitization. In contrast to the VAS ratings, behaviorally assessed pain perception did not differ between test and control trials after winning as well as after losing in the placebo condition (Figure 2 D; interaction ‘outcome × trial type’, *F*(1, 1040) = 2.53, *p* = 0.112).

### Levodopa increases endogenous pain modulation by active relief, naltrexone has no influence on the modulation

We next examined whether endogenous modulation of pain perception within the wheel of fortune game was affected by a levodopa and naltrexone.

#### Manipulation check: successful blinding of drug conditions

After the intake of *levodopa*, one participant reported a weak feeling of nausea and headaches at the end of the experimental session. In 32 out of 83 experimental sessions subjects reported tiredness at the end of the session. However, the frequency did not significantly differ between drugs (*χ*^2^ (2) = 2.17, *p* = 0.337). No other side effects were reported. To ensure that participants were kept blinded throughout the testing, they were asked to report at the end of each testing session whether they thought they received levodopa, naltrexone, placebo, or did not know. In 43 out of 83 sessions that were included in the analysis (52%), participants reported that they did not know which drug they received. In 12 out of 28 sessions (43%), participants were correct in assuming that they had ingested the placebo, in 6 out of 27 sessions (22%) levodopa, and in 2 out of 28 sessions (7%) naltrexone. The amount of correct assumptions differed between drug (*χ*^2^ (2) = 7.70, *p* = 0.021). However, post-hoc tests revealed that neither in the levodopa nor in the naltrexone condition participants guessed the correct pharmacological manipulation above chance level (*p’s* > 0.997), indicating that blinding was successful.

#### Ratings of perceived pain

As in the placebo condition, participants rated the thermal stimulation as less intense after active relief winning in the wheel of fortune task, and as more intense after receiving phasic pain increases (‘losing’) compared to the respective passive control condition under levodopa as well as naltrexone (Figure 2 B & 2 C).

Moreover, the effect of active relief or increases on pain modulation was differentially modulated by the drugs (interaction ‘drug × outcome’, *F(2*, 1587.30) = 4.52, *p* = 0.011). Specifically, the effect of active relief on perception was larger in the levodopa condition compared to the placebo condition (post-hoc comparison *p* = 0.007; Figure 3 A). No such difference was found for the naltrexone condition (*p* = 0.252). Endogenous modulation did not significantly differ between the levodopa and the naltrexone condition (*p* = 0.368). Endogenous pain facilitation induced by actively receiving pain increases assessed with VAS ratings did not significantly differ between any drug conditions (all post-hoc comparisons *p’s* > 0.591).

**Figure 3:**
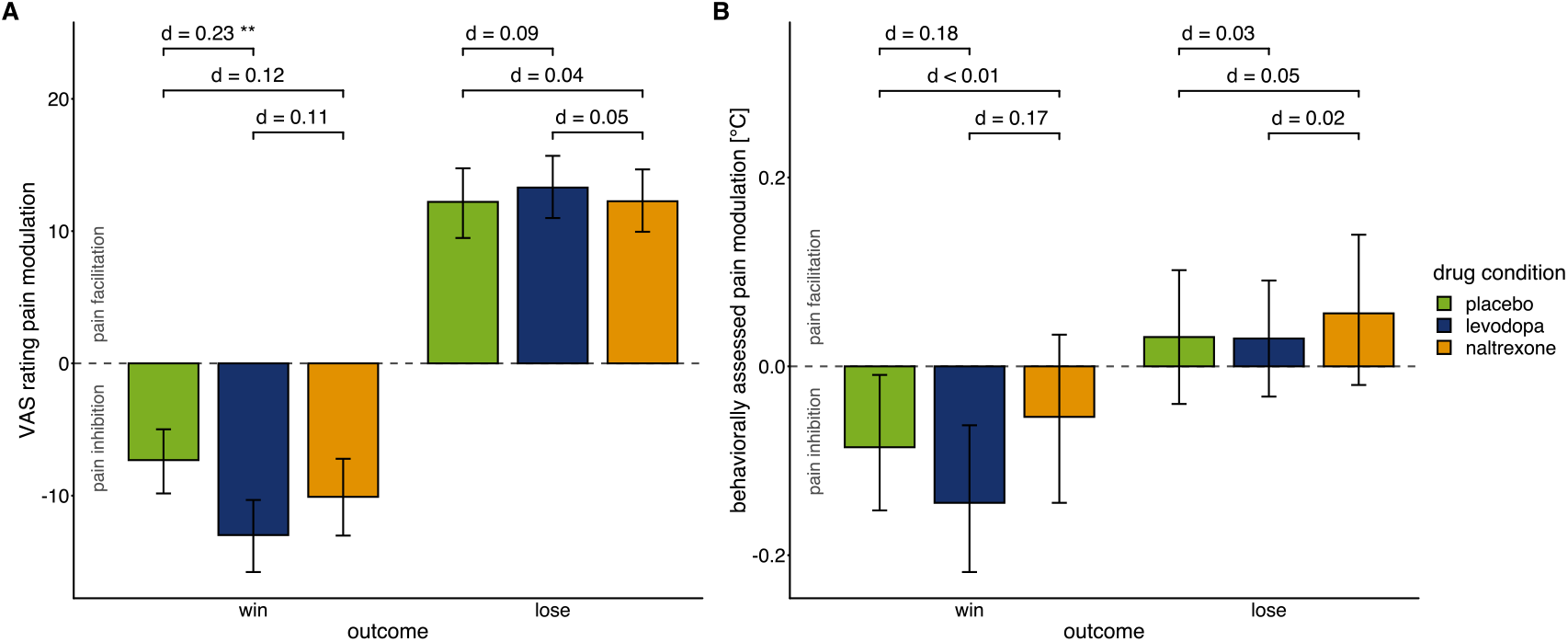
Effects of drug manipulation on endogenous pain modulation assessed by VAS ratings of pain intensity (A) and behaviorally assessed pain perception (B) after winning and losing in the wheel of fortune game, respectively (placebo: n = 28, levodopa: n = 27, naltrexone: n = 28). Error bars show 95% confidence interval of the mean. d indicates Cohen’s d as standardized effect-size of estimated effects. While the temporal order of sessions did affect pain modulation (figure supplement 1), measures of pain sensitivity, that were not experimentally manipulated (figure supplement 2), and measures of mood (figure supplement 3) did not significantly differ between drug conditions.

Endogenous pain inhibition under placebo and levodopa showed a high positive correlation (*r* = 0.591, *p* = 0.001). This correlation suggests that levodopa linearly increased endogenous pain-inhibitory effects of actively winning relief in the game dependent on endogenous pain inhibition mechanism in the placebo condition. In summary, the levodopa results show that the enhanced of relief perception during active controllability is dopamine-sensitive.

#### Behaviorally assessed pain perception

In contrast to the placebo condition, participants showed less behaviorally assessed sensitization in active compared to passive trials when obtaining pain relief under levodopa (Figure 2 E) consistent with an extension of pain-inhibitory effects of winning pain relief through to the underlying tonic pain stimulus. Under naltrexone, test and control trials did not significantly differ in the behaviorally assessed pain perception (Figure 2 F) as for the placebo condition. Across drugs, behaviorally assessed pain modulation did not significantly differ between placebo, levodopa, and naltrexone (interaction ‘drug × outcome’: *F*(2, 1592.73) = 1. 87, *p* = 0.154; Figure 3 B).

### Levodopa and naltrexone influence relief reinforcement learning in the wheel of fortune task

To investigate whether pain relief gained in active relief seeking was associated with an impact on choice related to reinforcement learning, one of the 2 choices in the wheel of fortune was associated with a fixed 75% chance of winning pain relief (*choice_high prob_*) while the other choice only had a 25% chance to win pain relief (*choice_low prob_*). Participants were not informed of these probabilities in advance. We tested if the proportion of choices of the more rewarding option was higher in the last two out of five blocks of four test trials each of the game, when the subjects already had the chance to explore and learn the different outcome probabilities.

Participants selected the color of the wheel of fortune associated with a higher likelihood for winning relief in 64% (*SD* = 28%) of in the placebo condition, consistent with a reinforcement learning effect. Thus, participants chose the color associated with the higher likelihood for winning above chance (*χ*^2^(1) = 6.64, *p* = 0.010) on a group level, indicating successful learning.

However, participants’ performance significantly differed between the placebo and the drug conditions (main effect of ‘drug’: *χ*^2^(2) = 11.89, *p* = 0.003). In contrast to the placebo condition (post-hoc comparison *p* < 0.001), under levodopa and under naltrexone participants’ choices did not significantly differ from chance (post-hoc comparisons *p’s* > 0.759). Correspondingly, post-hoc comparisons show that choice behavior significantly differed in the placebo compared to the levodopa condition (*p* = 0.015) and compared to the naltrexone condition (post-hoc comparison *p* = 0.004), while choices did not significantly differ between levodopa and naltrexone (post-hoc comparison *p* = 0.915; Figure 4). This shows that both, dopamine and opioids, may have an influence on relief-related learning and choice.

**Figure 4:**
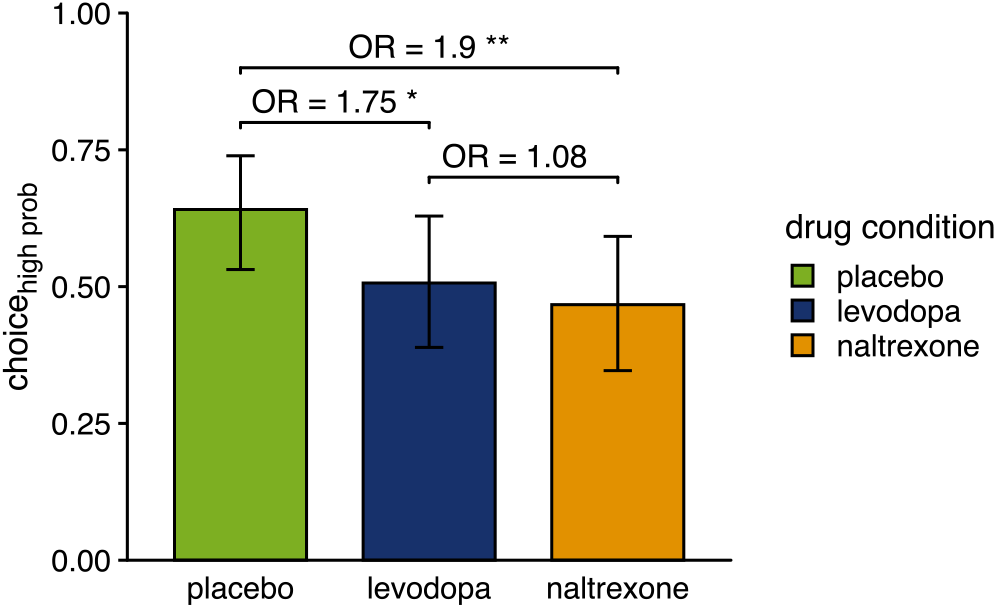
Proportion of choices of the color associated with a higher chance of winning pain relief (placebo: n = 28, levodopa: n = 27, naltrexone: n = 28). OR indicates odds ratios as effect size of estimated effects between drugs. \**p*<0.05, ** *p*<0.01.

In an additional exit interview at the end of each session, participants were asked whether they believed that one color of the wheel was associated with a higher chance of winning pain relief. The proportion of participants who reported this color correctly was not above chance (binomial test: *p’s* > 0.5; placebo: 50%, levodopa: 37%, naltrexone: 39.3%). Nevertheless, participants’ belief whether one color of the wheel of fortune task was associated with a higher chance of winning or not significantly influenced their choices (*p* < 0.001) and this influence on choices, and thus on learning, depended on the drug condition (interaction ‘drug × belief’: *F*(2) = 6.91, *p* = 0.032). Group effects of successful learning, i.e. selecting the color with a higher chance of winning, were driven by participants who were able to report this association (*p*(*choice*_*high prob*_|*correct belief*) = 0.737, *p*(*choice*_*high prob*_|*false or no belief*) = 0.545; post-hoc comparison: *p* = 0.007) under placebo and naltrexone (*p’s* < 0.001) but not under levodopa (*p* = 0.922). This suggests that successful decision-making was at least partly dependent on explicit contingency awareness.

### Unpredictability and endogenous pain modulation

We next tested whether outcome unpredictability was associated with endogenous pain modulation, and whether this prediction differed between drugs. Prediction errors describe the difference between an expected and a received outcome for positive (here pain relief) as well as negative outcomes (here phasic pain increases) (Glimcher, 2011; Schultz, 2016), and thus capture a measure of unpredictability or surprise, that determines how much learned values need to be updated. To obtain estimates for such prediction errors, we fit different reward learning models, with a drift diffusion process as the choice rule to participants’ choice and reaction time data. The best predictive accuracy was found for model 4 that used an individually scaled outcome sensitivity, and a sigmoid function to map expected values for the two choices to the drift rate of the diffusion process (Table 2; see *Methods and Materials*, section *Estimation of prediction errors and their role in endogenous pain modulation* for details on parametrization of reward learning models).

**Table 1:**
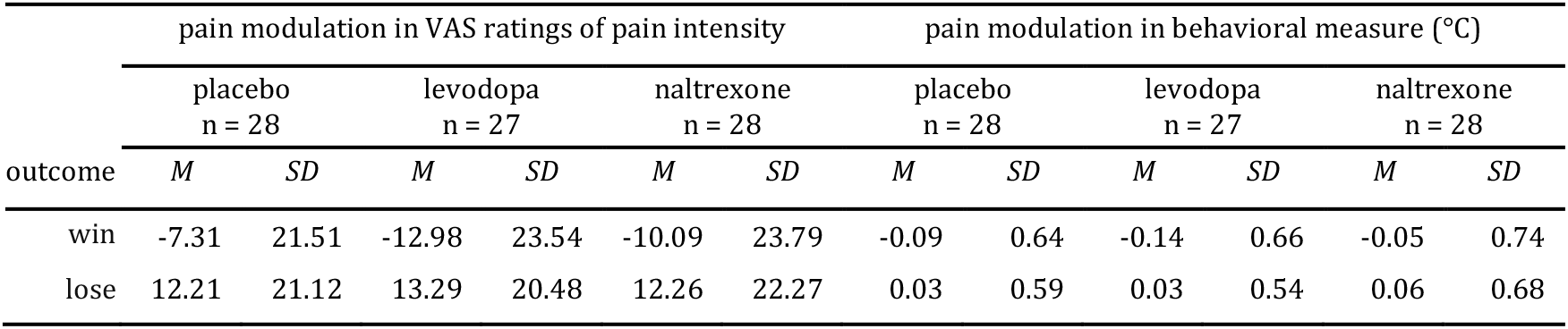
Means and standard deviation of means for pain modulation in VAS ratings of perceived intensity and the behaviorally assessed pain perception (negative values indicate pain inhibition; positive values indicate pain facilitation).

**Table 2:**
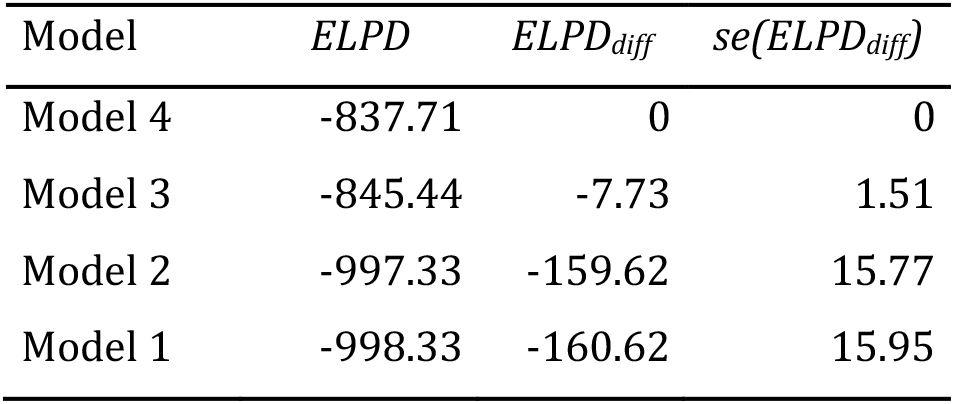
Model comparison. Models are ordered by their expected log pointwise predictive density (*ELPD*). *ELPD_diff_*: difference to the *ELPD* of winning model 4. *se*(*ELPD_diff_*): standard error of the difference in *ELPD*.

Posterior predictive simulations from the best-fitting model appropriately describe the observed choices (Figure 5). However, none of the model parameters could exclusively explain the differences between levodopa and naltrexone compared to placebo: the 95% highest density intervals (HDI) for the difference between all group level parameters of the drug effect enclosed zero (see Figure 5, figure supplement 1). Among the parameters affecting value updating (positive (*η*_+_) and negative (*η*_−_) learning rate and outcome sensitivity (*ρ*)) only *η*_−_ showed marginally higher central tendency for naltrexone compared to placebo, indicating a higher learning rate for punishments, but the 95% HDI still enclosed zero. The parameters affecting the mapping of expected values to the drift rate (*ν*, *ν*_max_) as well the other parameters affecting the drift diffusion decision process (non-decision time *τ*, boundary separation *α*, and a-priori bias *β*) were comparable in the placebo, levodopa, and naltrexone conditions.

**Figure 5:**
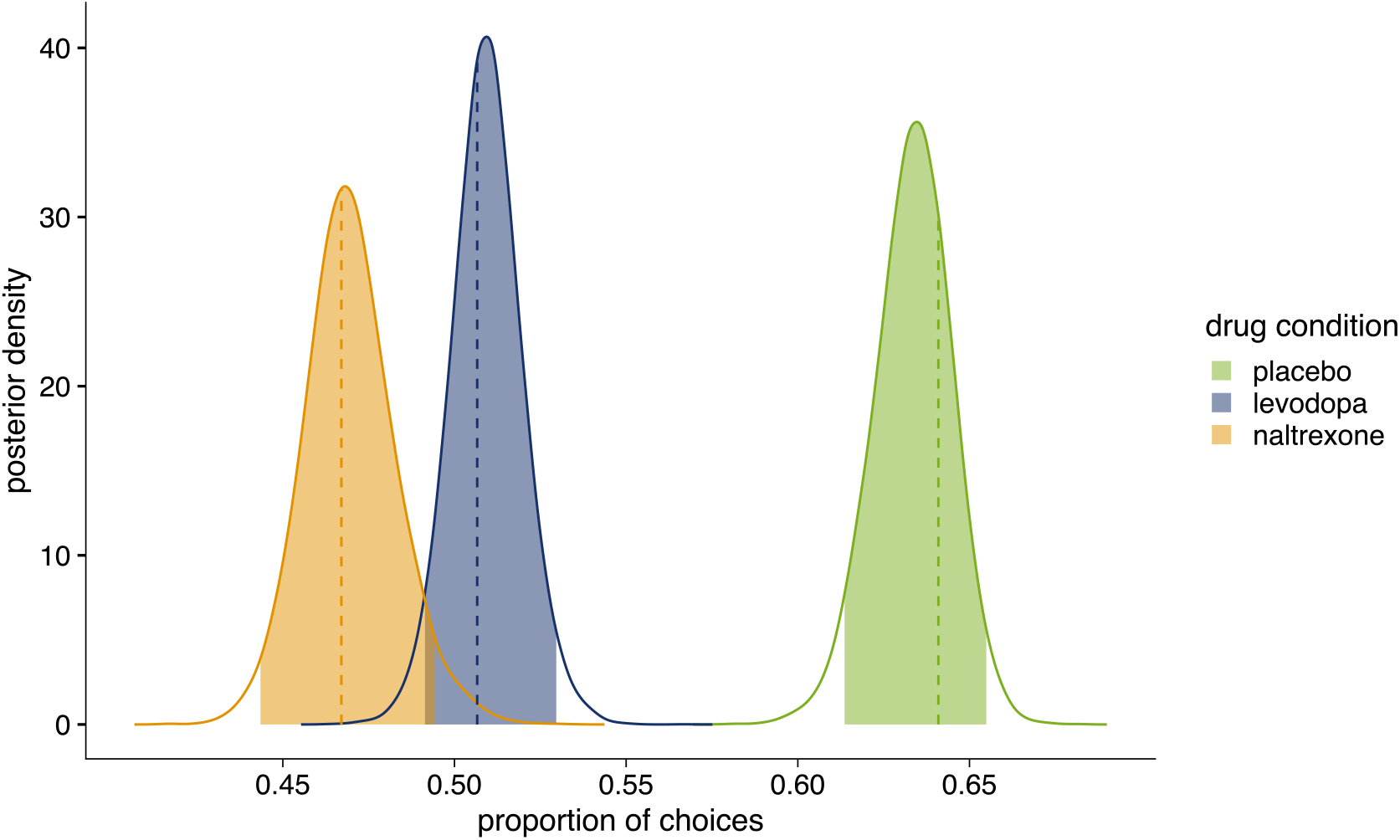
Posterior distribution of the proportion of choices in favor of *choice_high prob_* (placebo: n = 28, levodopa: n = 27, naltrexone: n = 28). Colored areas show 95% highest density interval (*HDI_95_*). Dashed lines indicate observed proportion of choices in favor of *choice_high prob_*. Placebo: *p*(*choice_high prob_*) = 0.641, *HDI_95_* = [0.614,0.655], posterior p-value (*pp*) = 0.320); levodopa: *p*(*choice_high prob_*) = 0.507, *HDI_95_* = [0.491,0.530], *pp* = 0.679; naltrexone: *p*(*choice_high prob_*) = 0.467, *HDI_95_* = [0.443,0.494], *pp* = 0.611. Figure supplement 1 shows comparison of drug conditions for each parameter of winning model 4.

Prediction errors estimated by using subject level parameters of the model showed a significant main effect for the prediction of endogenous pain modulation indicated by VAS ratings (*F*(1, 1600.3) = 452.9, *p* < 0.001). A negative estimate of the prediction error (*β_PE_* = −0.36) indicates that outcomes that are better than expected (positive prediction errors, which occur when receiving relief) were related to increased relief perception (pain inhibition). Conversely outcomes that are worse than expected (negative prediction errors, occurring with pain increases) were associated with increased pain facilitation (Figure 6). In other words, the more unexpected the relief, the greater the perception of that relief; and the more unexpected the pain increase, the greater the perception of that pain.

**Figure 6:**
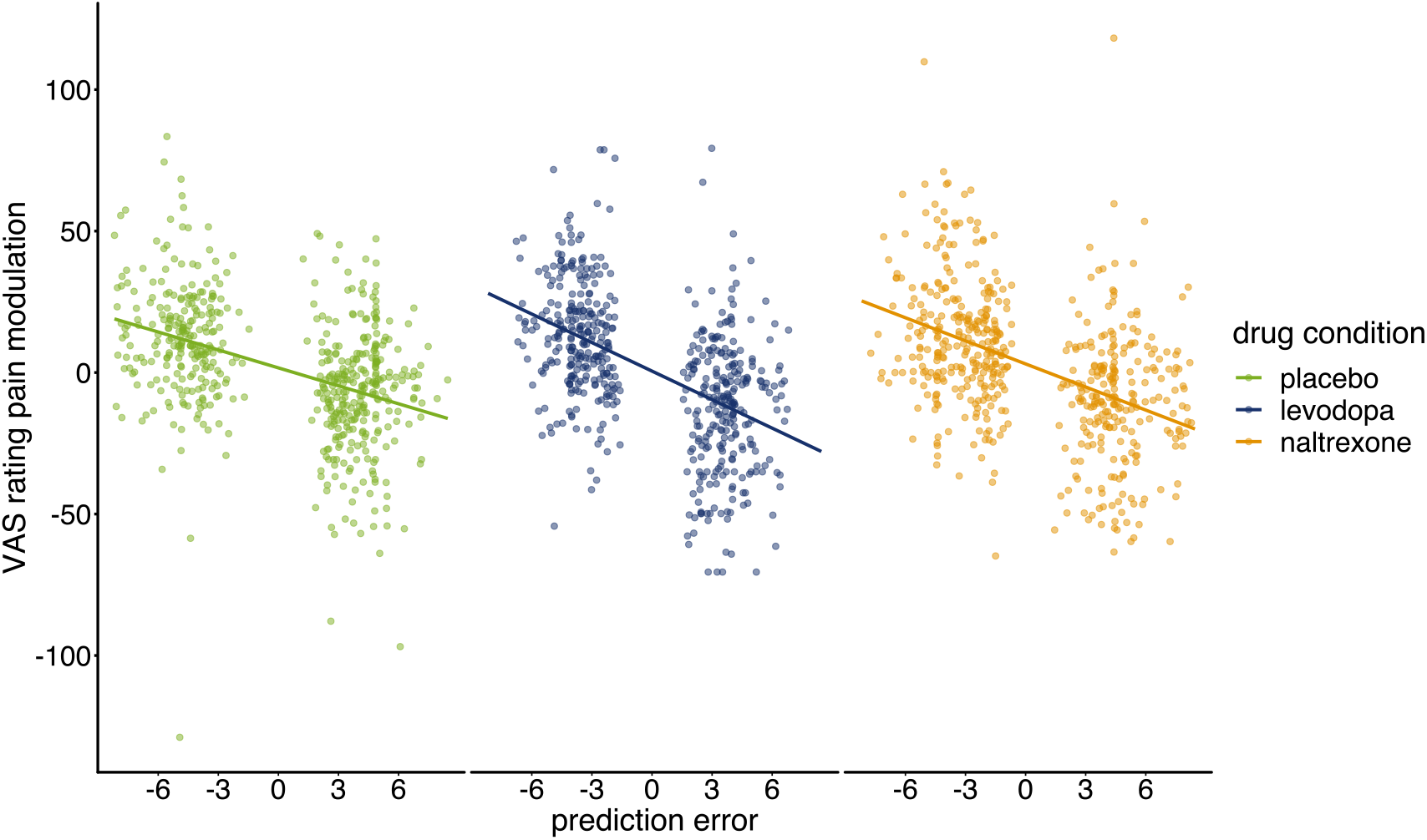
Pain modulation in VAS ratings predicted by prediction error for each condition (placebo: n = 28, levodopa: n = 27, naltrexone: n = 28). Regression lines indicate prediction from the mixed effects model with predictors ‘PE’, ‘drug’, and their interaction.

The effect of prediction errors on pain modulation showed a significant interaction with the drug condition (*F*(2, 1599.5) = 7.529, *p* < 0.001). Post-hoc analysis confirmed that the negative linear relationship significantly differed from zero for all conditions (*p’s* < 0.001), but this relationship was significantly stronger for levodopa compared to placebo (*p* < 0.001) with no significant differences for naltrexone compared to placebo (*p* = 0.083). Overall, this shows that relief is enhanced to unpredictability, and this effect is sensitive to dopamine.

Estimated prediction errors also showed a significant main effect for the prediction of behaviorally assessed pain modulation (*F*(1, 1602.1) = 9.00, *p* = 0.003), with a negative estimate (*β_PE_* = −0.06) suggesting that sensitization decreased with smaller prediction errors. No significant interaction with of prediction error with drug conditions was found for behaviorally assessed pain perception (interaction ‘PE × drug’: *F*(2, 1600.1) = 0.96, *p* = 0.384).

### Novelty seeking is linearly associated with increased endogenous pain modulation by pain relief under levodopa

Previous data suggest that endogenous pain inhibition induced by actively winning pain relief is associated with a novelty seeking personality trait: greater individual novelty seeking is associated with greater relief perception (pain inhibition) induced by winning pain relief (Becker et al., 2015). Replicating these results, we found here that endogenous pain modulation, assessed using self-reported pain intensity, induced by winning was correlated with participants’ scores on novelty seeking in the NISS questionnaire (Need Inventory of Sensation Seeking; Roth & Hammelstein, 2012; subscale ‘need for stimulation’ (NS); *r* = −0.412, *p* = 0.036). A similar association between novelty seeking and endogenous pain modulation was found in the levodopa condition (*r* = −0.551, *p* = 0.004). More importantly, the higher a participants’ novelty seeking score in the NISS questionnaire, the greater the levodopa-related endogenous pain modulation when winning compared to placebo (NISS NS: *r* = −0.483, *p* = 0.017, Figure 7). Pain modulation after losing was not associated with novelty seeking in placebo (*r* = 0.083, *p* = 0.687) and levodopa (*r* = −0.164, *p* = 0.433).

**Figure 7:**
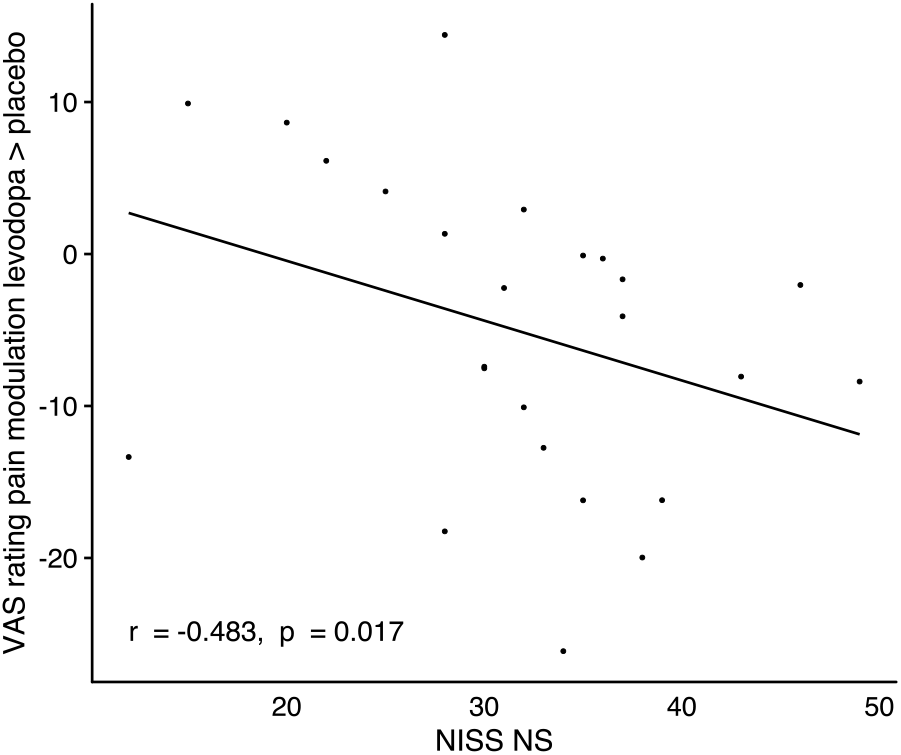
Correlation of changes in endogenous pain modulation induced by winning pain relief under levodopa compared to placebo with individuals’ scores on the ‘need for stimulation’ subscale of the NISS questionnaire, n = 24.

No significant correlations with NISS novelty seeking score were found for behaviorally assessed pain modulation in the placebo and levodopa conditions during pain relief or pain increase (|*r*|*’s* < 0.24, *p’s* > 0.266). Similarly, the difference in pain modulation during pain relief or pain increase between the levodopa and the placebo condition did also not correlate with novelty seeking (|*r*|’s < 0.22, *p’s* > 0.295).

## Discussion

The results show that i) the perception of relief is sensitive to endogenous modulation during motivated behavior, ii) this modulation scales with the informational content of the relief, being enhanced when relief is actively controllable, more unexpected, and especially in high trait novelty seeking individuals, iii) this information-specific modulation is sensitive to manipulation of dopamine signaling, with no evidence of a role of opioidergic signaling; iv) however both dopaminergic and opioidergic signaling have an influence on relief-seeking, which may be at least in part dissociable from relief perception. Overall, this shows that dopaminergic signaling is involved in a fundamental component of the endogenous modulation of pain relief.

Theories of the endogenous modulation of pain propose that one of the reasons that pain is modulated is to optimize motivational behavior, in terms of responding, learning, and decision-making (Fields, 2018; Seymour, 2019). That is, pain is increased in situations in which it has a more important role in shaping behavior – for instance when it directs a change in behavior (instrumentally controllable), when it is partly unpredictable (i.e. contains new information), and in otherwise dangerous contexts. This theory centralizes the functional role of pain as a signal for behavioral control i.e. concerned with the *prospective* control of behavior. In principle, this can be extended as a potential account for the modulation of relief, because the offset of pain is also important as a control signal for guiding behavior, one which occurs in the context of an ongoing noxious event, such as an injury of some sort. We have previously found preliminary evidence of this, by showing that relief perception is enhanced by active controllability (Becker et al., 2015). Here we intended to test this more precisely, by looking at the role of controllability as well as unpredictability, and also compare to the modulation of phasic increases in tonic pain.

We also set an additional prediction, in that we expected to find that modulation by information content would be greater in novelty-seeking individuals (Becker et al., 2015). This is because novelty seeking describes an explicit information-seeking tendency, in which new information is explored with the potential to lead to knowledge of better outcomes that can be exploited in the future (Wittmann, Daw, Seymour, & Dolan, 2008). This illustrates the common basis for intrinsic motivation for novelty and information-seeking for exploitable benefit, and hence we can predict that high trait novelty seekers might be more sensitive to information that occurs through relief outcomes.

Overall, all three predictions were borne out by the data: relief perception as measured by VAS ratings was enhanced by controllability, unpredictability and novelty-seeking tendency, consistent with the hypothesis that relief is sensitive to the exploitable information it carries. This provides the first clear formal framework for understanding a key component of relief perception. The principles for controllability and unpredictability also extended to increases in pain, consistent with the notion that increases in tonic pain act in a similar way to phasic pain operating from a pain-free baseline.

Both dopamine and opioids are implicated in relief processing, although their precise roles remain unclear. We found endogenous relief modulation here was modulated by enhanced dopamine availability induced by the intake of levodopa. Importantly, all three core aspects of informational-sensitivity were modulated by levodopa: active controllability, unpredictability, and association with novelty seeking. In contrast to our hypothesis, pharmacologically blocking opioid receptors using naltrexone did not modulate endogenous pain inhibition in the context of the task. The doses and methods used here are comparable to those used in other contexts which have identified opioidergic effects (Chelnokova et al., 2014; Eikemo, Biele, Willoch, Thomsen, & Leknes, 2017; King et al., 2013; Sirucek et al., 2021), suggesting that opioidergic effects on relief information are at least not substantial.

These findings also illustrate potential parallels with the previous observation of endogenous pain inhibition by extrinsic monetary reward co-occurring with experimental pain (Becker et al., 2013). In this context, monetary reward represents an independent and potentially competing incentive, and when this co-occurs with pain, it means that optimal responding may require suppression of pain responses, especially innate responses that could interfere with reward acquisition. In both cases, the common principle may be the active ‘decision’ by the pain system to tune incoming pain signals to optimize behavior. Note that the personality trait of novelty seeking has also been associated with enhanced dopaminergic activity due to lower midbrain (auto)receptor availability (Leyton et al., 2002; Savage et al., 2014; Zald et al., 2008), which further supports a general role for dopamine in information-sensitive behavior (Kakade & Dayan, 2002; Vellani, De Vries, Gaule, & Sharot, 2020).

The role of dopamine in pain relief in the context of reinforcement is supported by findings of increased dopamine release induced by pain relief in the Nucleus accumbens of rats (Navratilova et al., 2012; Xie et al., 2014). Dopamine release was related to the development of conditioned place preference that could be blocked by dopamine antagonists (Navratilova et al., 2012). Further, Navratilova, Xie, et al. (2015) showed that dopamine release in the Nucleus accumbens and conditioned place preference in response to pain relief depend on opioidergic signaling: both were blocked by opioid antagonism in the anterior cingulate cortex, an area encoding pain aversiveness. In humans, Sirucek et al. (2021) showed that perception of passively received pain relief is at least partly mediated by opioidergic neurotransmission. However, in that task, received pain relief did not carry behaviorally relevant information. Increased opioid activity in the anterior cingulate cortex has been shown to be associated with selectively decreased pain aversiveness with unaltered sensory pain components (Gomtsian et al., 2019; Maruyama et al., 2018; Navratilova et al., 2015). In contrast, the present study aimed at quantifying the effect of controllability on the relief perception, with these methods possibly not capturing the effects of opioid blockade on positive affective quality components of the relief experience. Overall, the finding that the modulation of pain relief was not modulated by naltrexone may suggest the possibility that a genuinely opioid-independent mechanism causes this type of pain inhibition.

One key difference in the current version of the wheel of fortune task, compared to the previous version described in (Becker et al., 2015), is that participants’ choices had a non-random association with outcomes i.e. this was a true instrumental (operant) contingency between actions and outcomes. This allowed us to assess a basic measure of learning – whether subjects are able to learn to select more frequently the option with the better (75% chance of relief) over worse (25% chance of relief) outcome. That both levodopa and naltrexone conditions were associated with a reduction of the frequency of choosing the better option, indicates that signals mediated by both neurotransmitters may be involved in choice. However, the data argue against a simple transposition of experienced relief (measured by VAS) into decision value, which for a stationary task such as this, should lead to more deterministic actions in the levodopa condition but no effect under naltrexone compared to placebo. The association of explicit contingency awareness and choice in our task illustrates the fact that multiple decision systems (‘model-based’ and ‘model-free’) might be involved in even simple instrumental tasks, and hence that more sophisticated task manipulations are needed to decompose these different components (Langdon, Sharpe, Schoenbaum, & Niv, 2018). However, our key finding is that there is at least a simple dissociation between the drug effects on experienced relief and decision-making.

Such dissociation may be due to differential involvement of dopamine and endogenous opioids in different yet interacting aspects of reward and punishment processing. Dopamine has been related to instrumental learning due to its prominent role in mediating reward and aversive prediction errors (Glimcher, 2011; Matsumoto & Hikosaka, 2009; Schultz, 2007, 2016). Correspondingly, effects of dopaminergic modulation on value-based decision making and brain activity related to reward prediction errors in the Nucleus accumbens have been reported (Pessiglione, Seymour, Flandin, Dolan, & Frith, 2006). On the other hand, impaired learning functions under dopaminergic medication are known from research in Parkinson’s disease (Breitenstein et al., 2006; Pizzagalli et al., 2008; Santesso et al., 2009; Vo, Seergobin, Morrow, & MacDonald, 2016) and have been attributed to dopamine overstimulation (Cools, Barker, Sahakian, & Robbins, 2001; Vaillancourt, Schonfeld, Kwak, Bohnen, & Seidler, 2013). Others argued that dopamine overstimulation does not impair learning of associations or reward expectations, but only the transfer to overt actions (choice behavior)(Beeler, 2012; Beeler, Daw, Frazier, & Zhuang, 2010). Accordingly, Kroemer et al. (2019) found reduced model-free control of choice behavior under levodopa (i.e. a decrease in direct reinforcement of actions by rewards) while both, neural reward prediction error signals and also model-based learning remained unaffected.

Given the involvement of multiple decision systems in our task a potential overstimulation with dopamine could have led here to choices not being guided by values learned from reinforcement. At the same time, dopamine has also been implicated in motivational aspects (incentive salience) of reward processing (Berridge et al., 2009; Smith et al., 2011; Tindell et al., 2005). Hence, dopamine may have increased motivational drive and related facilitation of pain modulation in the present task, while at the same time increased dopamine availability may have interfered with reinforcement learning. Opioids have been related to both, incentive salience and the hedonic value of rewards (Berridge et al., 2009; Meier, Eikemo, & Leknes, 2021). In humans, bidirectional manipulations have shown that opioid agonism increases while opioid antagonism decreases “wanting” (i.e. incentive salience) as well as “liking” of attractive faces (Chelnokova et al., 2014). The same mechanism was also shown for the effort to work for and the response bias for higher monetary rewards indicating that opioid manipulations affect motivation but also choice behavior (Eikemo et al., 2017). Such effects could explain here why the participants in this study did not develop a preference for choosing more frequently the option associated with a higher chance to win pain relief under naltrexone.

The data may have clinical implications. Reward learning has recently been shown to play a role in the transition of acute to chronic pain with a specific pattern of Nucleus accumbens activity in response to a cue predicting pain relief being predictive for chronification (Löffler et al., in press). This makes pain relief processing a potential leverage point for prevention strategies. Although levodopa or dopamine agonists are not generally used as analgesics in the clinical management of chronic pain, it may be that they could have a potential adjuvant role in management programs, for example when used in the context of rehabilitation strategies that aim to harness endogenous control mechanisms. It is also worth noting that Parkinson’s disease has a well-recognized association with chronic pain, beyond that which can be explained by motor effects, and in keeping with a potential core role for dopamine in the pathogenesis of chronic pain in some contexts (Beiske, Loge, Rønningen, & Svensson, 2009).

In summary, our study shows that dopamine has a core role in pain relief information processing, by which it modulates the way in which information tunes the modulation of pain to meet motivational demands.

## Materials and Methods

### Participants

Thirty healthy volunteers (16 female, 14 male; age: mean = 27.1 years; SD = 7.9 years) participated in this study. Exclusion criteria were present pain or pain conditions in the last 12 months, mental disorders, excessive gambling, substance abuse behaviors, alcohol consumption of 100 ml or more of alcohol per week, regular night shifts, or sleep disorders. Based on previous studies a medium effect size was expected (Becker et al., 2015). The a priori sample size calculation for an 80% chance to detect such an effect at a significance level of *α*=0.05 yielded a sample size of 28 participants (estimation performed using GPower version 3.1; (Faul, Erdfelder, Lang, & Buchner, 2007) for a repeated-measures ANOVA with within-subject factors). The study was approved by the Ethics Committee of the Medical Faculty Mannheim, Heidelberg University, and written informed consent was obtained from all participants prior to participation according to the revised Declaration of Helsinki (World Medical Association, 2013).

### Testing sessions

Each participant performed three testing sessions on separate days. Each session comprised a pharmacological intervention and a wheel of fortune game to assess modulation of reward-induced endogenous pain modulation by the interventions. Participants received in one session levodopa to transiently increase the availability of dopamine, in one session the opioid receptor antagonist naltrexone to block opioid receptors, and in one session a placebo for control. To ensure complete washout of the drugs, the testing sessions were separated by at least 2 days (plasma half-life for levodopa: 1.4 hrs (Nyholm et al., 2012); plasma half-life for naltrexone: 8 hrs (Wall, Brine, & Perez-Reyes, 1981)). After obtaining written consent in the first testing session, participants were familiarized with the thermal stimuli, the rating scale, and the wheel of fortune game to decrease unspecific effects of novelty and saliency. In each testing session the thermal pain threshold and pain tolerance were assessed prior to playing the wheel of fortune game to determine the stimulation intensities in the wheel of fortune game.

### Thermal stimulation

All heat stimuli were applied using a 25 × 50 mm contact thermode (SENSELab—MSA Thermotest, SOMEDIC Sales AB, Sweden). The baseline temperature was set to 30°C. Rise and fall rates of the temperature were set to 5°C/s. All thermal stimuli were applied to the inner forearm of participants’ non-dominant hand after sensitization of the skin using 0.075% topical capsaicin cream to allow for potent pain relief as reward and pain increase as punishment without the risk of skin damage (Becker et al., 2015; Gandhi, Becker, & Schweinhardt, 2013). By activating temperature-dependent TRPV1 (vanilloid transient receptor potential 1) ion channels capsaicin as the active ingredient of chili pepper induces heat sensitization (Holzer, 1991). To ensure that the entire area of thermal stimulation during the wheel of fortune game was sensitized the cream was applied to an area on the forearm exceeding the area of stimulation by about 1 cm on each side. After 20 min, the capsaicin cream was removed (Dirks, Petersen,& Dahl, 2003; Gandhi et al., 2013) and the thermode was applied. If participants reported the baseline temperature of the thermode (30°C) as painful because of the preceding sensitization this temperature was lowered until it was perceived as non-painful, which was needed in 8 out of 83 sessions (3 placebo sessions, 1 levodopa session, and 4 naltrexone sessions) that were finally entered into the analysis (see below). The temperature was decreased to 28°C (1 placebo session, 4 naltrexone sessions) or 26°C (1 placebo session, 1 levodopa session). The need to lower the baseline temperature was not significantly different between drug conditions (Fisher’s exact test, *p* = 0.52).

### Determination of stimulation intensities

Participants’ heat pain threshold and heat pain tolerance were assessed using the method of limits three times prior to the wheel of fortune game. The temperature of the thermode increased from baseline with 1°C/s. Participants were instructed to press the left button of a three-button computer mouse when the pain threshold was reached. The respective temperature was recorded while the temperature further increased. Participants were instructed to press the button again when the pain tolerance threshold was reached. The respective temperature was recorded and the temperature immediately returned to baseline. The arithmetic mean of the temperatures corresponding to the recorded pain threshold and tolerance in the three trials was used as an estimate of the individual heat pain threshold and heat pain tolerance, respectively.

After this threshold and tolerance assessment, an adjustment procedure resembling a staircase method was implemented to determine the stimulation intensities in the wheel of fortune game. Participants received heat stimuli of 20s duration and continuously rated the perceived intensity of these stimuli on a computerized visual analogue scale (VAS) ranging from “no sensation” (0) over “just painful” (100) to “most intense pain tolerable” (200) (Becker et al., 2013; Villemure et al., 2003) while the stimuli where presented. The temperature of the first trial was set to the mean of the previously determined pain threshold and tolerance. If the rating at the end of the stimulation was outside a range of 150±10 on the VAS, the temperature for the next trial was adjusted according to the difference to a target rating of 150. This adjustment was determined by multiplying the difference (150 - current rating) by 0.02 and adding the result in °C to the previous temperature. Further, temperature increases between trials were limited to a maximum of 0.5°C to avoid overshooting of ratings. The procedure was repeated until a rating between 140 and 160 on the VAS was achieved, aiming at a temperature perceived as moderately painful. The corresponding temperature was used as the stimulation intensity in the wheel of fortune game.

### Wheel of fortune game

A wheel of fortune game, adapted from a previously established version (Becker et al., 2015), was used to provide participants with the possibility of winning pain relief. The game comprised three types of trials: *test* trials, in which participants played the wheel of fortune game and received pain relief or pain increases according to the outcome of the game; *control* trials, in which participants did not play the game, but received pain relief or pain increases as in the test trials; and *neutral* trials, in which participant did not play the game and no pain relief or pain increases were implemented. A trial always started with an increase of the temperature to the previously determined tonic pain stimulation intensity. When the stimulation intensity was reached, participants were instructed to memorize the temperature perceived at this moment (Figure 1). After this memorization interval, participants were presented with a wheel of fortune display that was divided into three sections of equal size but different color.

In the *test* trials, participants were asked to select one of two colors (pink or blue) of the wheel by pressing a corresponding button (left or right) on the mouse. This started the wheel spinning (4.3 s) until it stopped on either the blue or pink section. When the wheel came to a stop and the pointer of the wheel indicated the color the participant had chosen, the stimulation temperature decreased with the aim to induce pain relief (win condition). If the pointer indicated the color the participant had not chosen, the temperature was increased (lose condition). In the *control* trials, participants had to press a black button unrelated to the sections of the wheel of fortune using the middle mouse button, after which the wheel started spinning as in the test trials. In contrast to the test trials, no pointer was displayed in the control trials and the wheel stopped at a random position. After the wheel came to a stop, the stimulation temperature decreased or increased, resembling the course of stimulation in the test trials, but without winning or losing. By this procedure, nociceptive input in test and control trials was kept the same, allowing to test specifically for endogenous pain modulation induced by winning and losing in the wheel of fortune game.

In *neutral* trials, participants had to press a black button, as in the control trials, after which the wheel also started spinning. In these neutral trials, the pointer of the wheel always landed the third color of the wheel (white), which could not be selected in test trials, and the stimulation temperature did not change. Neutral trials were used to estimate changes in pain perception occurring over the course of the experiment due to habituation or sensitization independent of the outcomes of the wheel of fortune game.

After the interval of the temperature change (in the test trials: outcome of the wheel), participants rated the perceived intensity of the current temperature using the same VAS as described above (Figure 1). After this rating, participants had to adjust the stimulation temperature themselves to match the temperature they had memorized at the beginning of the trial. This self-adjustment operationalizes a behavioral assessment of perceptual sensitization and habituation within one trial (Becker et al., 2015, 2011; Kleinböhl et al., 1999). Participants adjusted the temperature using the left and right button of the mouse to increase and decrease the stimulation temperature. Self-adjusted temperatures lower than the stimulation intensity at the beginning of the trial indicate sensitization across the trial, while higher temperatures indicate habituation. After this behavioral assessment, the stimulation temperature went back to baseline and after a short break (5 s) the next trial started.

In total, the wheel of fortune game comprised of 45 trials, split into five blocks. Each block consisted of 4 *test* and 4 *control* trials followed by one *neutral* trial. *Test* and *control* trials were presented in a predefined, pseudorandomized sequence. In contrast to the previous version of the wheel of fortune (Becker et al., 2015), the outcome of the wheel occurred with certain likelihood to allow for learning to optimize the outcomes of the wheel of fortune. One of the colors (pink or blue) was associated with a 75% chance of winning, while the other was associated with a 25% chance of winning (counterbalanced across participants and testing sessions). If participants did not select a color in the test trials, the neutral outcome (white) of the wheel was displayed and the temperature did not change. The temperature changes in the *control* trials (pain relief or increase) were matched to the outcomes of the *test* trials to ensure that the same number of pain relief and pain increase trials were presented in test and control trials.

Pain relief was implemented by a reduction of the stimulation intensity of 3°C and pain increase was implemented by a rise of 1°C. The magnitude of these temperature steps was determined and optimized in pilot experiments with the aim of inducing potent pain relief and pain increase without inducing ceiling and floor effects.

Although the main focus of the study was to test different effects on pain relief as implemented in win trials and their corresponding control trials with a decrease in nociceptive input, lose trials and their complementing control trials were crucial to the experimental design. First, for playing the game lose trials were an integral part because of the implemented likelihood for winning which necessarily needs to be accompanied by the chance of losing. Additionally, the risk of losing was thought to increase the participants’ engagement in the game, which in turn was expected to enhance the motivated state induced by playing the wheel of fortune game. Second, they allowed for testing whether pain modulation was driven by controllability or unspecific effects such as arousal and distraction in test compared to control trials of the wheel of fortune game (c.f. Becker et al., 2015).

All experimental procedures involving thermal stimulation were controlled by custom-programmed Presentation scripts (Presentation^®^ software, Version 17.0, http://www.neurobs.com) providing instructions and other visual cues on a computer screen in front of the participants.

### Pharmacological manipulations

Participants ingested in one testing session *levodopa*, in another *naltrexone*, and in another a placebo (microcrystalline cellulose), following a double-blind, cross-over design with counterbalanced order. Levodopa is an amino acid precursor of dopamine leading to a transient systemic increase of dopamine availability. To inhibit peripheral synthesis of dopamine from levodopa, the single dose of 150 mg levodopa (p.o.) was combined with 62.5 mg of a benserazide to prevent peripheral side effects such as nausea (Rinne et al., 1975). Naltrexone is an opioid receptor antagonist with predominant receptor binding affinity at μ-opioid receptors together with a lower binding affinity at κ-opioid receptors and a much lower affinity at δ-opioid receptors (Raynor et al., 1994). Participants received a single dose of 50 mg naltrexone (p.o.) that has been shown to induce more than 90% receptor blockade (Weerts et al., 2013).

After drug intake, a waiting period of one hour started. This waiting time was chosen based on peak plasma concentrations of levodopa and naltrexone at approximately 1 h to 1.5 h after ingestion (Nyholm et al., 2012; Wall et al., 1981). At the end of each testing session, participants indicated whether they thought that they had received the placebo or one of the drugs (response alternatives: ‘placebo’, ‘levodopa’, ‘naltrexone’, or ‘don’t know’) to test for potential unblinding.

### Questionnaire and exit interview

Novelty seeking as personality trait was assessed using the Need Inventory of Sensation Seeking (NISS; Roth and Hammelstein, 2012). The NISS consists of the sub-scales *Need for Stimulation* (NS) and *Avoidance of Rest* (AR). We used the NS subscale as a measure for novelty seeking as it reflects the “need for novelty and intensity” (Roth & Hammelstein, 2012). Before playing the wheel of fortune game the affective state of subjects was assessed using computerized versions of the Self-Assessment Manikin (SAM; Bradley & Lang, 1994; Lang, 1980) and a German version (Krohne, Egloff, Kohlmann, & Tausch, 1996) of the Positive And Negative Affect Scale (PANAS; Watson, Clark, & Tellegen, 1988). At the of each session, an exit interview was performed, asking for the following information: (1) which drug participants believed to have ingested; (2) if participants believed that choosing one of the two colors was associated with a higher chance to win pain relief; (3) whether participants perceived a difference between test and control trials; (4) whether participants had the impression that the stimulation temperature at the beginning of each trial varied across trials; (5) whether participants had problems indicating their perception on the VAS scale; and (6) whether participants had problems readjusting the initial temperature. Participants gave first yes/no answers and then were asked to specify their answers using open-ended questions.

### Statistical analysis

For the statistical analysis, 2 participants were excluded, one participant due to the failure to comply with experimental procedures and one due to technical failure of the equipment. For one additional participant, data of one session (levodopa) are missing due to drop-out. Thirty-two out of 3735 single trials of all the remaining sessions were not recorded due to technical failures. In 42 trials, participants did not press a button within the respective interval in the wheel of fortune game. These trials were excluded from the analyses. Note that the NISS questionnaire was missing for two additional subjects due to initial issues at the beginning of the data collection.

To test if blinding was successful we fit a mixed-effects logistic regression with the subjects’ assumption on the ingested drug (as reported in the exit interview, see above) being correct as dependent variable. We used ‘drug’ as a fixed factor and to account for repeated measures we modelled a random intercept for each subject. Post-hoc general linear hypothesis tests were used to compare estimated proportions of correct assumptions against chance.

To confirm that the manipulation of the motivated state (test vs. control trials) of the participants by playing the wheel of fortune game did induce pain modulation as intended in each session, we analyzed the VAS ratings and the behavioral pain measure as outcome measures separately for each session with ‘trial type’ and ‘outcome’ as well as their interaction as fixed effects. To account for the repeated measures design we modelled a random intercept for each participant and a random slope for outcome of the wheel within each participant.

To obtain an estimate for endogenous pain modulation in each test trial, we subtracted the mean value of all control trials of either the pain relief or the pain increase trials from the value of the winning or losing test trials separately for each session for both the VAS ratings and the behavioral pain measure. Using these differences, negative values indicate pain inhibition and positive values indicate pain facilitation. Estimates for pain modulation were analyzed using linear mixed model procedures with the fixed factors ‘drug’ (levodopa, naltrexone, placebo), ‘outcome’ (win, lose), ‘order’ of sessions (1, 2, 3), and their interaction separately for ratings and behaviorally assessed pain perception as dependent variables. The factor ‘session number’ was added to control for effects of temporal order independent of the drug manipulation that was found to influence pain modulation (see Figure 3, figure supplement 1). Other factors such as baseline pain perception or mood did not affect pain modulation and were not included in further analysis (see Figure 3, figure supplement 2 and 3). To account for the repeated measures design we modelled a random intercept for each participant and a random slope for outcome within each participant.

Unbeknown to the subjects, one of the colors in the wheel of fortune was associated with a higher chance to win pain relief. To test whether participants learned to select this color from the implemented reward contingencies we looked at choice behavior in the last 2 blocks of trials only. In this latter phase of the task subjects already had the chance to explore differences in outcomes associated with their choices and were thought show exploitation if they had learned about the contingency. We fit a mixed-effects logistic regression with the subjects’ choices as dependent variable. For a single session we fit an intercept only model where the intercept represents the group level estimate for the probability to choose the color associated with a higher chance of winning pain relief (*choice_high prob_*). Drug was used as an additional within-subject factor when testing for differences among levodopa, naltrexone, and placebo. To account for repeated measures, we modelled a random intercept for each subject. To assess the effect of the subjects’ belief about which color was associated with a higher chance to win pain relief (as reported in the exit interview, see above) we added the factor ‘belief’ (either “correct belief” or “false or no belief”) to this model.

To test whether endogenous pain modulation due to winning pain relief was related to participants’ personality trait of novelty seeking, pain modulation represented by the differences between test and control trials in the wheel of fortune in VAS ratings and the behaviorally assessed pain perception of the placebo and the levodopa condition were correlated with the NISS NS scores. To test further whether increases in pain modulation induced by levodopa were associated with novelty seeking, differences in pain modulation between the levodopa and placebo session were also correlated with the NISS NS scores. Before calculating these correlations, multivariate outliers were tested using a chi-square test on the squared Mahalanobis distance using an *a* of 0.025 (Filzmoser, 2016), leading to the exclusion of one value for the correlation with the difference of pain modulation between the levodopa and placebo session.

The significance level was set to 5% for all analyses. All statistical analyses were performed using statistical computing software R version 3.5.3 (R Core Team, 2019). Mixed model analyses were performed using the *lme4* package (Bates, Mächler, Bolker, & Walker, 2015). All linear mixed models were estimated using restricted maximum likelihood. Kenward-Roger correction as implemented in the *lmerTest* package (Kuznetsova, Brockhoff, & Christensen, 2017) was used to calculate test statistics and degrees of freedom to account for the sample size. For general linear mixed-effects models Wald *χ*^2^ was calculated using *car* package (Fox, John & Weisberg, 2011). Post-hoc tests and effect sizes were calculated on estimated marginal means using the *emmeans* package (Lenth, 2020) where appropriate. Tukey adjustment was used to account for multiple comparisons in post-hoc tests.

### Estimation of prediction errors and their role in endogenous pain modulation

To analyze how mechanisms of instrumental learning contribute to the observed choice behavior and how this related to reward-induced pain modulation we fitted reinforcement learning (RL) models to participants’ choices in test trials of the wheel of fortune game. Such models were initially formulated for associative learning (Bush & Mosteller, 1951; Rescorla & Wagner, 1972) and adapted for instrumental learning (Sutton & Barto, 1998). RL models assume that actions are chosen based on the expected outcome. Learning is described as the adaptation of expectations based on experiences. Thus, learning is driven by the discrepancy between a present expectation and the obtained outcome, namely the *prediction error*. The speed of adaption of the expectation is described by the *learning rate*, which defines the exponential decay of the influence of previous outcomes on the currently present expectation. For trial-by-trial instrumental learning paradigms, the update of the expectation of an outcome related to a given action (in the present study: choice in the wheel of fortune) is operationalized by calculating the expected value *Q* of a choice as follows:

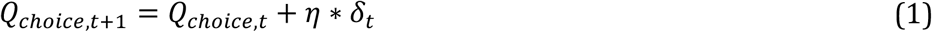

where *Q_choice_* is the reward expectation for a given choice, *t* denotes the trial, *η* is the learning rate, and *δ_t_* is the prediction error in trial *t*. The learning rate *η* determines the speed of adaption; the higher *η* the more is the expectation influenced by recent compared to former experiences. Since updating of expectations has been shown to differ dependent on the sign of the prediction error (Fontanesi, Gluth, Spektor, & Rieskamp, 2019; Gershman, 2015; Pedersen, Frank, & Biele, 2017), we modelled independent learning rates for positive (*η*_+_) and negative (*η*_−_) prediction errors:

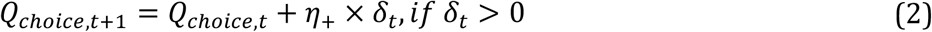

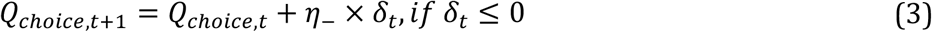

The prediction error as the difference between the actual and the expected outcome in trial *t* is formulated as follows:

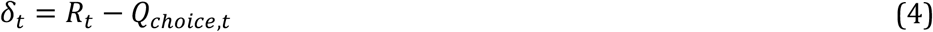

with *R_t_* as the outcome of the choice in trial *t*.

In the wheel of fortune game, outcomes were implemented as changes in stimulation intensities. Accordingly, *R_t_* was positive (+1) for temperature decreases in win trials or negative (−1) for temperature increases in lose trials. The formula shown above assumes a constant outcome sensitivity. To capture potential modulation of the outcome sensitivity, we implemented a scaled outcome sensitivity so that the reward in trial *t* was multiplied by an individual scaling factor *ρ* yielding a scaled prediction error:

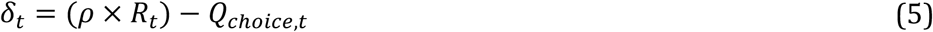

Q values were initiated to zero and calculated separately for choices of the color associated with a higher chance to win pain relief (*Q_high prob_*) and choices of the color associated with a lower chance to win pain relief (*Q_low prob_*).

While RL models traditionally used a softmax choice rule (Daw & Doya, 2006; Luce, 1959), recent studies on value-based decision making have implemented variants of the drift diffusion model (Ratcliff, 1978; Ratcliff & Rouder, 1998) to map expected values to choices (Fontanesi et al., 2019; Pedersen et al., 2017; Peters & D’Esposito, 2020). The drift diffusion model describes decisions as accumulation of noisy evidence for two choice options until a predefined threshold, representing either of the two options, is reached. Such drift diffusion models take response times (*RT*) of decisions into account and model mathematically cognitive processes underlying the decision process. Figure 8 depicts such a decision process. The range between the decision boundaries is represented by the boundary separation parameter *α*. Higher values of *α* lead to slower but more accurate decisions, that is, *α* represents the speed vs. accuracy tradeoff. The position of the starting point *z* between the boundaries is determined by *a priori* biases *β* toward one of the two options. This parameter *β* represents the relative distance of *z* between the boundaries. It can range from 0 to 1 where a value of 0.5 indicates no bias, values below indicate a bias for the lower choice and values above 0.5 a bias for the upper choice. The non-decision time *τ* describes time needed for processes that are unrelated to the decision process (e.g. stimulus processing). Correspondingly, the reaction time is defined as *RT* = *τ* + *decision time*. Acquisition of evidence starts from the starting point *z* at time *τ* as a random walk. The slope of this random walk is determined by the drift rate *v* and a decision is made when either the upper or lower boundary is reached. Higher drift rates result in faster and more accurate decisions. The probability of the *RT* when choosing option *x* can then be calculated using the Wiener first-passage time distribution (Ratcliff, 1978):

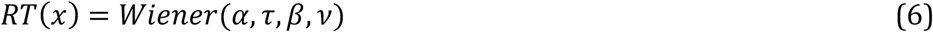

where *Wiener*() returns the probability that *x* is chosen with the observed RT.

**Figure 8:**
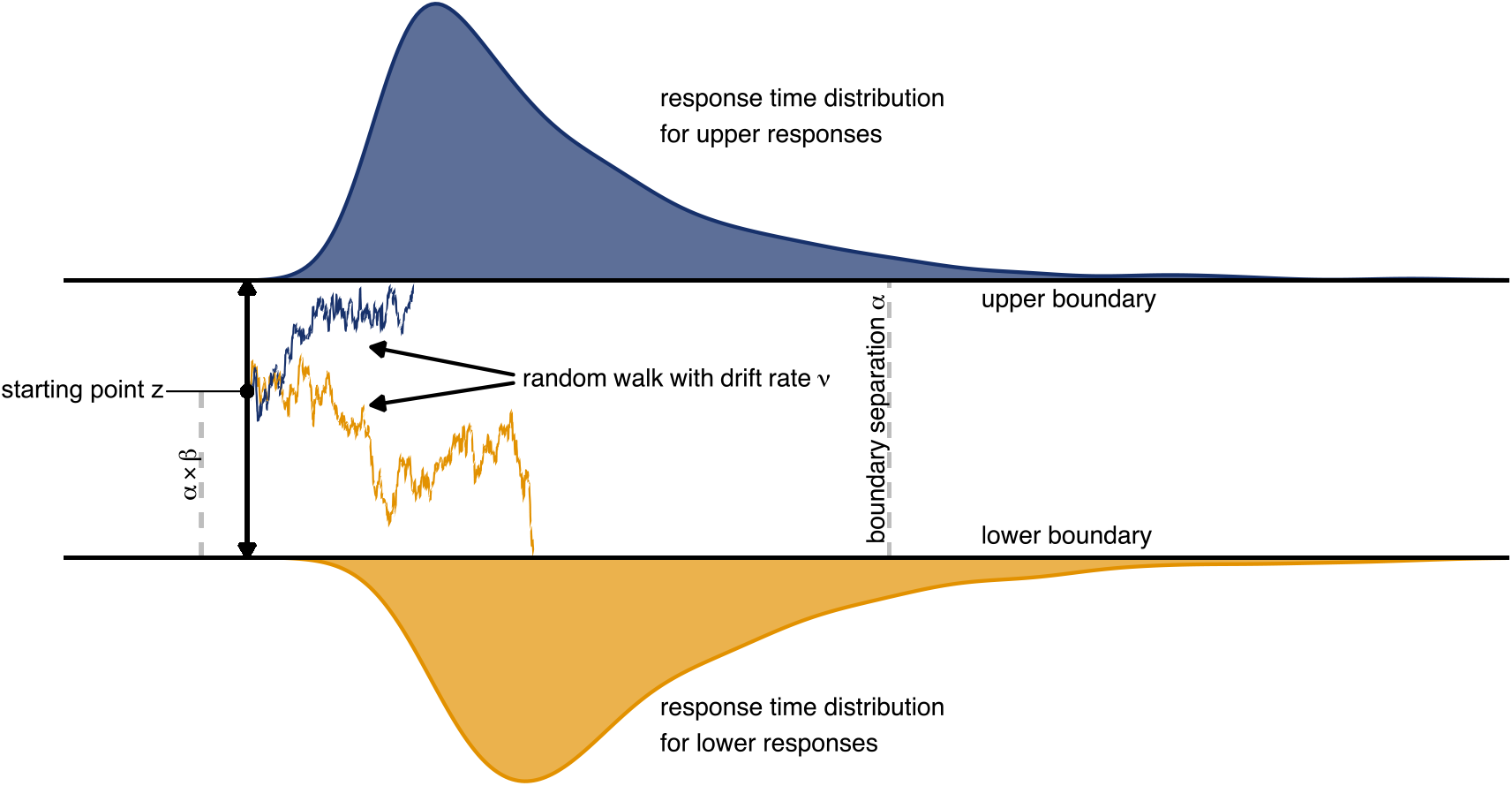
Schematic depiction of the drift diffusion model. Accumulation of evidence starts at point *z* which is defined by the a-priori bias *β* and the boundary separation *α*. Noisy evidence is integrated over time (represented by sample paths in blue and orange, for upper and lower boundary choices, respectively).

Most variants of reward learning models that use the drift diffusion process as a choice rule replace the constant drift rate by an individually scaled difference of expected values for the both options (Fontanesi et al., 2019; Pedersen et al., 2017; Peters & D’Esposito, 2020). Thus, the drift rate *v_t_*, varies across trials as a function of the difference between expected values of the two choice options that in the wheel of fortune corresponded to *Q_high prob_* and *Q_low prob_*, respectively. We implemented a linear mapping of the difference in expected values like Pedersen al. (2017) where this difference is multiplied by the scaling factor *v*:

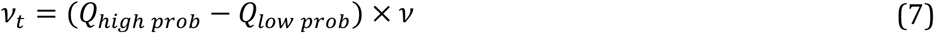

As an alternative scaling method we implemented a non-linear function as suggested by Fontanesi et al. (2019) in which the scaled difference in expected values is mapped to the drift rate using a sigmoid function, which more closely resembles the non-linear mapping of the softmax function:

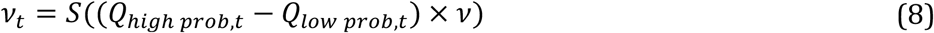

where *S*(*x*) is defined as:

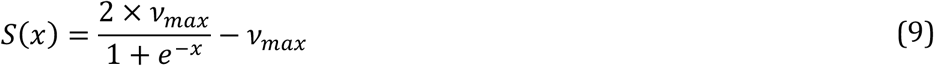

With that, ±*v*_max_ defines the upper and lower limit of the drift rate, respectively, while the shape or slope of the sigmoid function depends on the scaled difference of expected values.

In summary, we combined different parameterizations of the outcome sensitivity (static or scaled) and the mapping of expected values to the drift rate (linear or sigmoidal) into different models (Table 3).

**Table 3:**
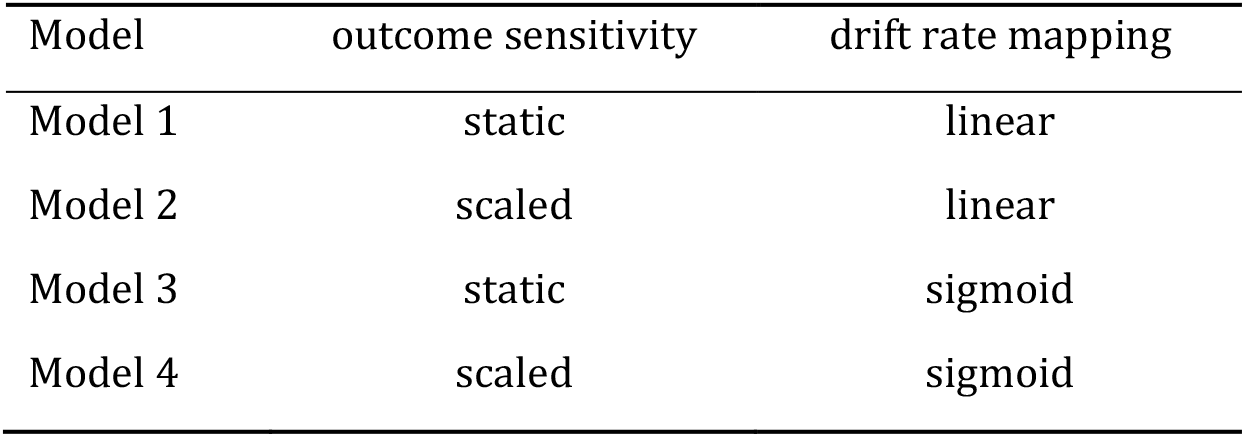
Model specification. Models 1-4 were defined using different combinations of parameters for reward sensitivity and the mapping of expected values to the drift rate. A ‘static’ reward sensitivity means that pain increase and pain decrease were defined as −1 and 1, respectively (see Equation 4). A ‘scaled’ outcome sensitivity means that pain decrease was defined as –*ρ* and pain decrease as *ρ* (see Equation 5). A ‘linear’ drift rate mapping means that the drift rate *v_t_* for each trial was defined as the difference of expected values multiplied by *v* (see Equation 7). A sigmoid mapping of the drift rate means that *v_t_* was defined by a sigmoid function bounded at ±*v*_max_. (see Equation 8 und Equation 9). All models included two learning rates (*η*_+_, *η*_−_), the non-decision time *τ*, the boundary separation *α*, and the a priori bias *β*.

| Model | outcome sensitivity | drift rate mapping |
| --- | --- | --- |
| Model 1 | static | linear |
| Model 2 | scaled | linear |
| Model 3 | static | sigmoid |
| Model 4 | scaled | sigmoid |

We used hierarchical Bayesian modeling to fit the reward learning models to the choices of the participants in the test trials. Hierarchical models estimate group and individual parameters simultaneously to mutually inform and constrain each other, which yields reliable estimates for both, individual and group level parameters (Gelman, Carlin, Stern, & Rubin, 2013; Kruschke, 2014). Posterior distributions of the parameters were estimated using Hamiltonian Monte Carlo sampling with a No-U-Turn sampler as implemented in the probabilistic language Stan (Carpenter et al., 2017) via its *R* interface *rstan* (Stan Development Team, 2020). For each model parameter, we included a global intercept and the main effect of drug (levodopa, naltrexone, placebo). Both, intercept and main effect were allowed to vary for each participant and we modelled a correlation of individual terms for the drug effect across participants to account for repeated measures. We used a non-centered parameterization to reduce dependency between group and individual level parameters (Betancourt & Girolami, 2015). Therefore, both intercept and drug effect were defined by their location (group level effect), scale, and error (individual effects) distributions. A logistic transformation was applied to the learning rate (*η*_+_, *η*_−_) and a-priori bias (*β*) parameters to restrict values to the range of [0, 1]. The location parameters for the intercept of the learning rate were given standard normal priors 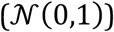 and the scale of these parameters were given half-normal priors 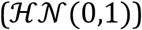. The location of the drug effect on learning rate parameters were also given standard normal priors while the scale was given a half-normal prior of 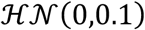 to prevent allocation of high prior density at the edges of the range after logistic transformation, resulting in an almost flat prior. The location parameter for the intercept of the a-priori bias was given a normal prior of 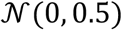 and the scale was given a half-normal prior 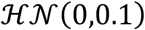. The location parameter for the drug effect was given a normal prior of 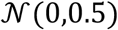 and the scale was given a half-normal prior of 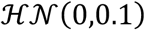. To ensure that the non-decision time (*τ*) was bounded to be lower than the reaction time the parameter was equivalently transformed to the range [0,1] and multiplied with each subject’s individual minimum reaction time in a given session. Priors were the same as for the learning rate, i.e. yielding a flat prior after transformation. We used an exponential transformation to constrain the reward sensitivity parameter (*ρ*), the boundary separation (*α*), drift rate scaling factor (*v*), and the boundary of the drift rate (*v*_max_) to be greater than 0. The location of the global intercept was given a normal prior of 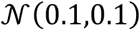 for the reward sensitivity, a normal prior of 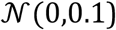 for the boundary separation, a normal prior of 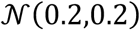 for the drift rate, and a normal prior of 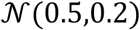 for the drift rate boundary. For the exponentially transformed parameters the scale of the global intercept was given a half-normal prior of 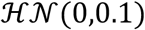, the location of the drug effect was given a normal prior of 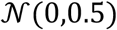, and the scale of the drug effect was given a half-normal prior of 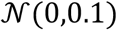. Individual effects for the intercept as well as for the drug effect were all given standard normal priors. The correlation matrix of individual drug-level effects for each parameter was given a LKJ prior (Lewandowski, Kurowicka, & Joe, 2009) of 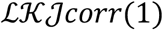. All models were run on four chains with 4000 samples each. The first 1000 iterations were discarded as warm-up samples for each chain. The convergence of chains was confirmed by the potential scale reduction factor 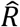.

The fitted models were compared for their best predictive accuracy using *K*-fold cross-validation (Vehtari, Gelman,& Gabry, 2017). For the cross-validation, we split data into *k* = 10 subsets with each subset containing data of 2-3 participants and calculated the expected log pointwise predictive density (*ELPD*) based on simulations for each hold-out set *y_k_* using parameters estimated from re-fitting the model to the training data set *y*_(−*k*)_. We calculated *ELPDs*, their differences, and the standard error of the differences using the *R* package *loo* (Vehtari et al., 2020). A higher *ELPD* indicates a better predictive accuracy. Such a better predictive accuracy was assumed if the difference in *ELPD* (*ELPD_diff_*) for two models was at least 2 times the standard error of that difference (*se*(*ELPD_diff_*)).

For the best fitting model, we performed posterior predictive checks by simulating replicated data sets from posterior draws. As the test statistic for the posterior predictive check we examined the proportion of choices in favor of the option associated with a higher chance to win pain relief (*choice_high prob_*) in the last 2 blocks of the wheel of fortune game and compared the proportions observed in this data to the distribution of proportions found in the simulated data sets.

From the best fitting model, we used group level estimates for the main effect of ‘drug’ to compare model parameters between drug conditions using the 95% highest density interval (HDI) of the difference of their posterior distributions.

The means of individual parameter posterior distributions were used to estimate prediction errors for single trials. To test whether these prediction errors predict endogenous pain modulation induced by the wheel of fortune task, we used linear mixed models with the fixed factors ‘prediction error’ and ‘drug’, and their interaction. A random intercept for each subject was included to account for repeated measures. Separate models for VAS ratings and behaviorally assessed pain perception as dependent variables were calculated.

## Acknowledgments

This work was funded by a Postdoctoral Fellowship for Leading Early Career Researchers from the Baden-Württemberg Foundation awarded to SB. Further, SB was supported by the Olympia Morata Program of the Heidelberg University and a PRIMA grant (PR00P1_179697/1) of the Swiss National Science Foundation. HF was supported by grants of the Deutsche Forschungsgemeinschaft (SFB1158 B03 and B07; Fl 156/41-1). BS is funded by Wellcome, Versus Arthritis (21537) and IITP (MSIT 2019-0-01371).

## Legends supplement figures

**Figure 3, figure supplement 1:**
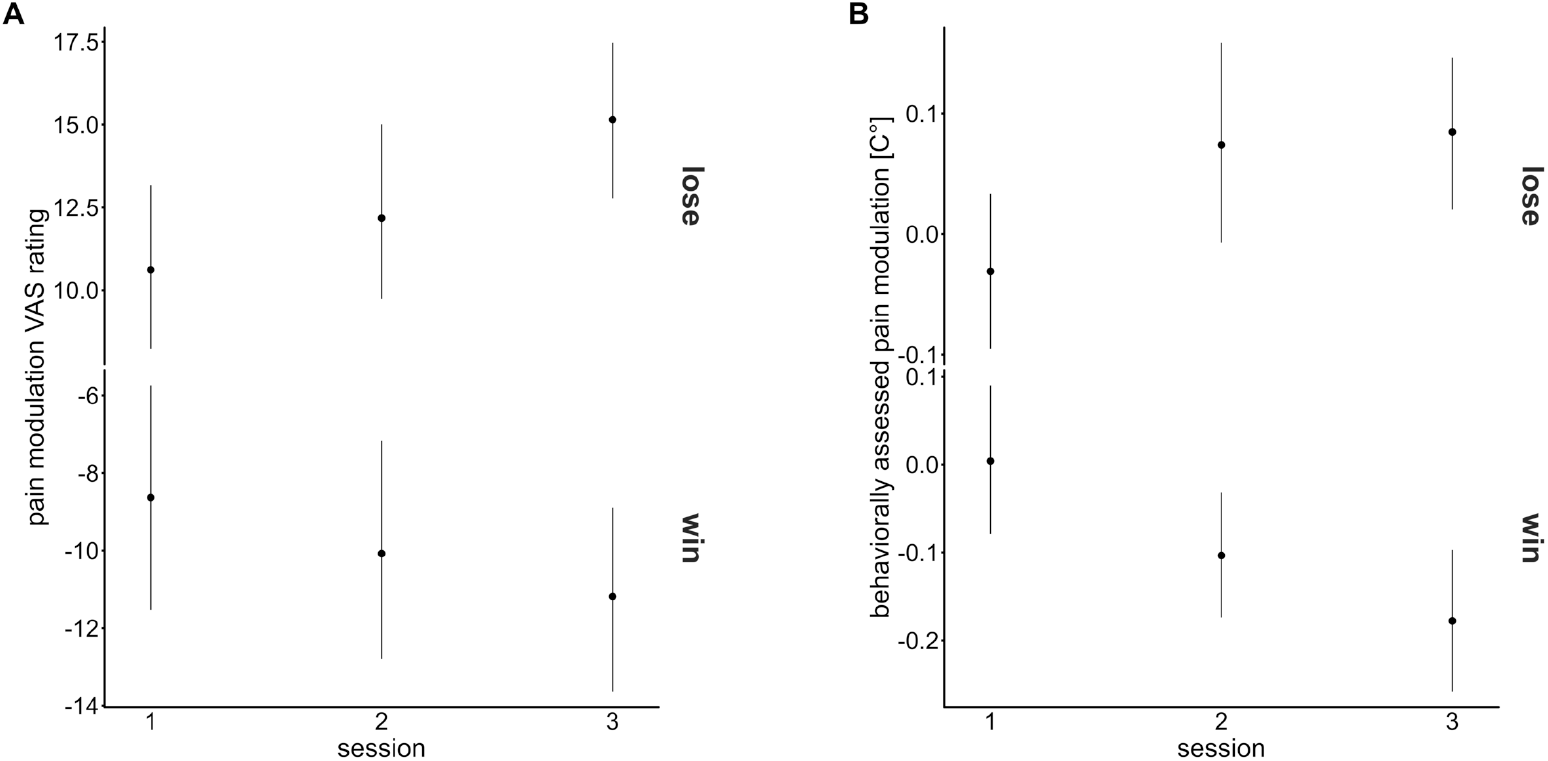
Means and 95% confidence intervals of the mean for pain modulation in (A) VAS ratings and (B) behaviorally assessed pain modulation for each testing session. Mixed-effects models testing whether the temporal order of the testing sessions, independent of the order of the application of the drugs, had an effect on pain modulation in win and lose trials of the wheel of fortune did not show a main effect of ‘session number’ (pain modulation VAS ratings: *F*(2, 1593.70) =1.28, *p* = 0.279; behaviorally assessed pain modulation: *F*(2, 1599.84) = 0.86, *p* = 0.425) but point to a differential effect of temporal order for win and lose outcomes (interaction ‘outcome × session number’: pain modulation VAS ratings: *F*(2, 1593.77) = 3.00, *p* = 0.050; behaviorally assessed pain perception: *F*(2, 1597.27) = 7.94, *p* < 0.001). Hence, temporal order was included as an additional main effect when testing the effect of ‘drug’ on pain modulation.

**Figure 3, figure supplement 2:**
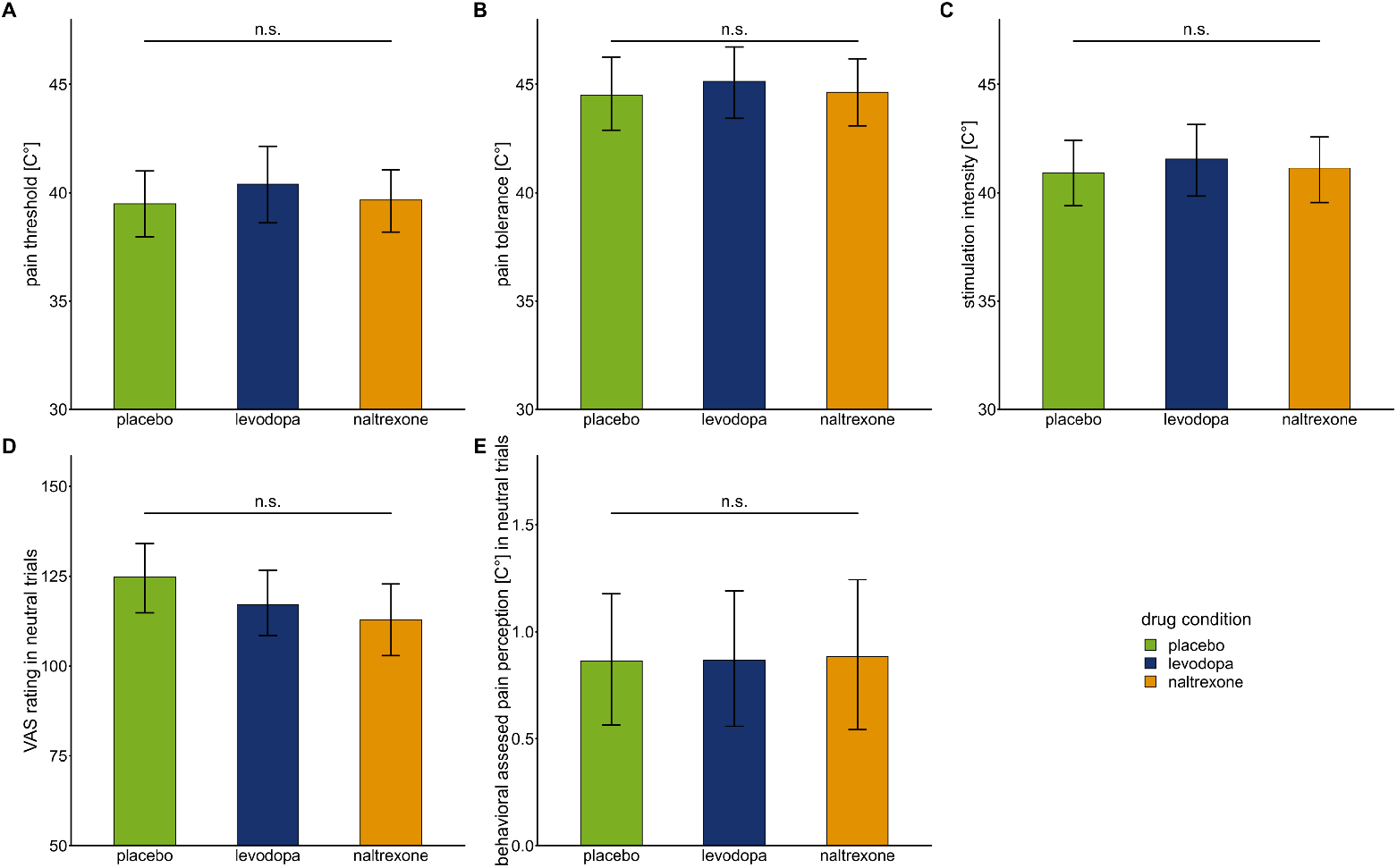
Bars show means and error bars 95% confidence intervals of the mean for (A) pain threshold, (B) pain tolerance, (C) stimulation intensity, (D) VAS ratings in neutral trials (in which participants did not play the game and the temperature stayed constant), and (E) behaviorally assessed pain perception in neutral trials for each drug condition. Mixed-effects models using drug condition (placebo: n = 28, levodopa: n = 27, naltrexone: n = 28) to predict measures of baseline pain sensitivity showed no significant main effect for ‘drug’: pain threshold: *F*(2,53.21) = 0.64, *p*=0.529; pain tolerance: *F*(2,53.18) = 0.31, *p* = 0.736; stimulation intensity: *F*(2,53.2) = 0.30, *p* = 0.745. Mixed-effects models using drug condition to predict VAS ratings and behaviorally assessed pain perception in the neutral condition of the wheel of fortune task showed no significant main effect for ‘drug’: VAS ratings: *F*(2,53.31) = 2.12, *p* = 0.131; behaviorally assessed pain perception: *F*(2,53.13) = 0.01, *p* = 0.990.

**Figure 3, figure supplement 3:**
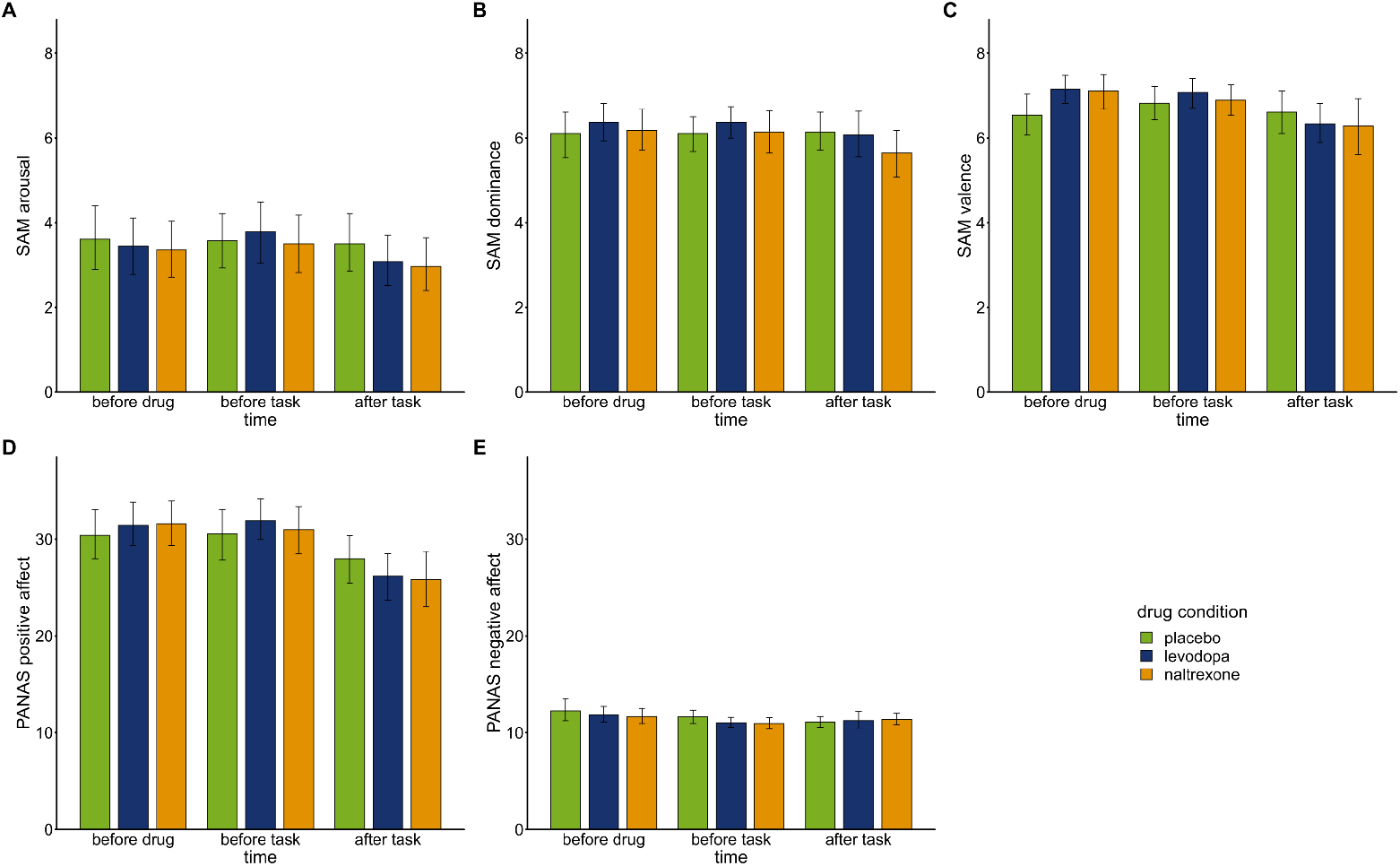
Mood was assessed over the course of each experimental session before drug intake, before playing the wheel of fortune game, and after playing the game using computerized versions of the Self-Assessment Manikin (SAM; Bradley & Lang, 1994; Lang, 1980) and a German version (Krohne, Egloff, Kohlmann, & Tausch, 1996) of the Positive And Negative Affect Scale (PANAS; Watson, Clark, & Tellegen, 1988). Bars show means and error bars 95% confidence intervals of the mean for SAM subscale (A) arousal, (B) dominance, (C) valence, and PANAS subscales (D) positive affect, and (E) negative affect at each time point. To test whether drug conditions (placebo: n = 28, levodopa: n = 27, naltrexone: n = 28) differentially affected mood we fit separate mixed-effects models predicting subscales of SAM and PANAS by ‘drug’, ‘time’, and their interaction. SAM ratings for arousal, dominance, and valence did not show any significant main effects of ‘drug’ (arousal: *F*(2,213.2) = 1.56, *p*=0.214); dominance: *F*(2,213.29) = 1.03, *p* = 0.359; valence: *F*(2,213.41) = 0.74, *p* = 0.479) nor significant interactions for ‘drug × time’ (arousal: *F*(4,213.0) = 0.69, *p* = 0.599; dominance: *F*(4,213.00) = 0.88, *p* = 0.4771; valence: *F*(4,213.00) = 2.28, *p* = 0.062). Participants’ positive affect assed with the PANAS did not show a significant main effect of ‘drug’ (*F*(2,213.25 = 0.05, *p* = 0.954) nor a significant interaction of ‘drug x time’ (*F*(2, 213.00) = 1.60, *p* = 0.176). Similarly, negative affect assessed with the PANAS did not show a significant main effect of ‘drug’ (*F*(2, 213.51) = 0.93, *p* = 0.376) nor a significant interaction of ‘drug × time’ (*F*(2, 213.00) = 0.79, *p* = 0.533).

**Figure 5, figure supplement 1:**
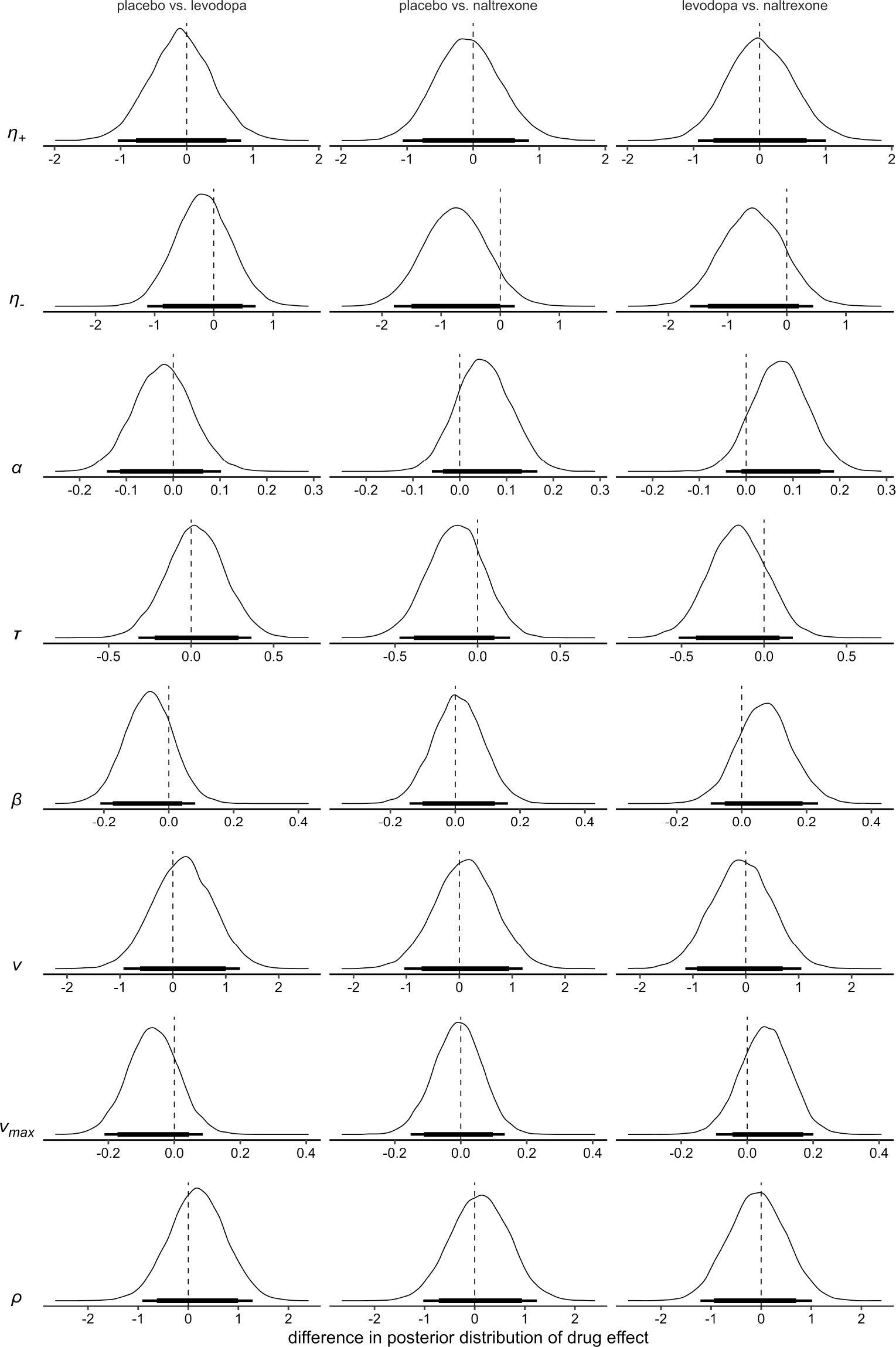
Differences of the posterior distributions of group level parameters for the main effect of drug in model 4. Thick black bars indicate the 85% HDI, thin bars indicate the 95% HDI. *η*_+_: learning rate for positive prediction errors; *η*_−_: learning rate for negative prediction errors; *α*: boundary separation; *τ*: non-decision time; *β*: a-priori bias; *v*: drift-rate scale factor; *v_max_* drift-rate boundary; *ρ*: outcome sensitivity.

## References

Bannister, K. (2019). Descending pain modulation: influence and impact. Current Opinion in Physiology, 11, 62–66. https://doi.org/10.1016/j.cophys.2019.06.004

Barbano, M. F., & Cador, M. (2006). Differential regulation of the consummatory, motivational and anticipatory aspects of feeding behavior by dopaminergic and opioidergic drugs. Neuropsychopharmacology, 31(7), 1371–1381. https://doi.org/10.1038/sj.npp.1300908

Barbano, M. F., & Cador, M. (2007). Opioids for hedonic experience and dopamine to get ready for it. Psychopharmacology, 191(3), 497–506. https://doi.org/10.1007/s00213-006-0521-1

Bates, D., Mächler, M., Bolker, B. M., & Walker, S. C. (2015). Fitting linear mixed-effects models using lme4. Journal of Statistical Software, 67(1), 1–48. https://doi.org/10.18637/jss.v067.i01

Becker, S., Gandhi, W., Elfassy, N. M., & Schweinhardt, P. (2013). The role of dopamine in the perceptual modulation of nociceptive stimuli by monetary wins or losses. European Journal of Neuroscience, 38(7), 3080–3088. https://doi.org/10.1111/ejn.12303

Becker, S., Gandhi, W., Kwan, S., Ahmed, A. K., & Schweinhardt, P. (2015). Doubling your payoff: Winning pain relief engages endogenous pain inhibition. ENeuro, 2(4), 1–11. https://doi.org/10.1523/ENEURO.0029-15.2015

Becker, S., Kleinböhl, D., Baus, D., & Hölzl, R. (2011). Operant learning of perceptual sensitization and habituation is impaired in fibromyalgia patients with and without irritable bowel syndrome. Pain, 152(6), 1408–1417. https://doi.org/10.1016/j.pain.2011.02.027

Beeler, J. (2012). Thorndike’s Law 2.0: Dopamine and the Regulation of Thrift. Frontiers in Neuroscience, 6. https://doi.org/10.3389/fnins.2012.00116

Beeler, J., Daw, N., Frazier, C., & Zhuang, X. (2010). Tonic Dopamine Modulates Exploitation of Reward Learning. Frontiers in Behavioral Neuroscience, 4. https://doi.org/10.3389/fnbeh.2010.00170

Beiske, A. G., Loge, J. H., Rønningen, A., & Svensson, E. (2009). Pain in Parkinson’s disease: prevalence and characteristics. PAIN®, 141(1-2), 173–177.

Benedetti, F. (1996). The opposite effects of the opiate antagonist naloxone and the cholecystokinin antagonist proglumide on placebo analgesia. Pain, 64(3), 535–543. https://doi.org/10.1016/0304-3959(95)00179-4

Berridge, K. C., Robinson, T. E., & Aldridge, J. W. (2009). Dissecting components of reward: “liking”, “wanting”, and learning. Current Opinion in Pharmacology, 9(1), 65–73. https://doi.org/10.1016/j.coph.2008.12.014

Betancourt, M., & Girolami, M. (2015). Hamiltonian Monte Carlo for Hierarchical Models. Current Trends in Bayesian Methodology with Applications, 79–101. https://doi.org/10.1201/b18502-5

Bradley, M. M., & Lang, P. J. (1994). Measuring emotion: The self-assessment manikin and the semantic differential. Journal of Behavior Therapy and Experimental Psychiatry, 25(1), 49–59. https://doi.org/10.1016/0005-7916(94)90063-9

Breitenstein, C., Korsukewitz, C., Flöel, A., Kretzschmar, T., Diederich, K., & Knecht, S. (2006). Tonic dopaminergic stimulation impairs associative learning in healthy subjects. Neuropsychopharmacology, 31(11), 2552–2564. https://doi.org/10.1038/sj.npp.1301167

Bush, R. R., & Mosteller, F. (1951). A mathematical model for simple learning. Psychological Review, 58(5), 313.

Carpenter, B., Gelman, A., Hoffman, M. D., Lee, D., Goodrich, B., Betancourt, M., … Riddell, A. (2017). Stan: A probabilistic programming language. Journal of Statistical Software, 76(1). https://doi.org/10.18637/jss.v076.i01

Chelnokova, O., Laeng, B., Eikemo, M., Riegels, J., Løseth, G., Maurud, H., … Leknes, S. (2014). Rewards of beauty: the opioid system mediates social motivation in humans. Molecular Psychiatry, 19(7), 746–747. https://doi.org/10.1038/mp.2014.1

Cools, R., Barker, R. A., Sahakian, B. J., & Robbins, T. W. (2001). Enhanced or Impaired Cognitive Function in Parkinson’s Disease as a Function of Dopaminergic Medication and Task Demands. Cerebral Cortex, 11(12), 1136–1143. https://doi.org/10.1093/cercor/11.12.1136

Daw, N. D., & Doya, K. (2006). The computational neurobiology of learning and reward. Current Opinion in Neurobiology, 16(2), 199–204. https://doi.org/10.1016/j.conb.2006.03.006

Dirks, J., Petersen, K. L., & Dahl, J. B. (2003). The heat/capsaicin sensitization model: A methodologic study. Journal of Pain, 4(3), 122–128. https://doi.org/10.1054/jpai.2003.10

Eikemo, M., Biele, G., Willoch, F., Thomsen, L., & Leknes, S. (2017). Opioid Modulation of Value-Based Decision-Making in Healthy Humans. Neuropsychopharmacology, 42(9), 1833–1840. https://doi.org/10.1038/npp.2017.58

Eippert, F., Bingel, U., Schoell, E. D., Yacubian, J., Klinger, R., Lorenz, J., & Büchel, C. (2009). Activation of the Opioidergic Descending Pain Control System Underlies Placebo Analgesia. Neuron, 63(4), 533–543. https://doi.org/10.1016/j.neuron.2009.07.014

Faul, F., Erdfelder, E., Lang, A.-G., & Buchner, A. (2007). A Flexible statistical power analysis program for the social, behavioral, and biomedical sciences. Behavior Research Methods, 39(2), 175–191. https://doi.org/10.3758/BF03193146

Fields, H. L. (2006). A motivation-decision model of pain : The role of opioids. In Proceedings of the 11th world congress on pain (pp. 449–59).

Fields, H. L. (2007). Understanding How Opioids Contribute to Reward and Analgesia. Regional Anesthesia and Pain Medicine, 32(3), 242–246. https://doi.org/10.1016/j.rapm.2007.01.001

Fields, H. L. (2018). How expectations influence pain. PAIN, 159(9), S3–S10. https://doi.org/10.1097/j.pain.0000000000001272

Filzmoser, P. (2016). Identification of Multivariate Outliers: A Performance Study. Austrian Journal of Statistics, 34(2), 127–138. https://doi.org/10.17713/ajs.v34i2.406

Fontanesi, L., Gluth, S., Spektor, M. S., & Rieskamp, J. (2019). A reinforcement learning diffusion decision model for value-based decisions. Psychonomic Bulletin and Review, 26(4), 1099–1121. https://doi.org/10.3758/s13423-018-1554-2

Fox, John & Weisberg, S. (2011). An R Companion to Applied Regression (Third Edit). Thousand Oaks, CA: Sage. Retrieved from http://cran.r-project.org/web/packages/car/citation.html

Gandhi, W., Becker, S., & Schweinhardt, P. (2013). Pain increases motivational drive to obtain reward, but does not affect associated hedonic responses: A behavioural study in healthy volunteers. European Journal of Pain, 17(7), 1093–1103. https://doi.org/10.1002/j.1532-2149.2012.00281.x

Gelman, A., Carlin, J. B. B., Stern, H. S. S., & Rubin, D. B. B. (2013). Bayesian Data Analysis, Third Edition (Texts in Statistical Science). Book, (February), 675. https://doi.org/10.1007/s13398-014-0173-7.2

Gershman, S. J. (2015). Do learning rates adapt to the distribution of rewards? Psychonomic Bulletin and Review, 22(5), 1320–1327. https://doi.org/10.3758/s13423-014-0790-3

Glimcher, P. W. (2011). Understanding dopamine and reinforcement learning: The dopamine reward prediction error hypothesis. Proceedings of the National Academy of Sciences of the United States of America, 188(SUPPL. 3), 15647–15654. https://doi.org/10.1073/pnas.1014269108

Gomtsian, L., Bannister, K., Eyde, N., Roble, D., Dickenson, A. H., Porreca, F., & Navratilova, E. (2019). Morphine effects within the rodent anterior cingulate cortex and rostral ventromedial medulla reveal separable modulation of affective and sensory qualities of acute or chronic pain. Physiology & Behavior, 176(3), 139–148. https://doi.org/10.1016/j.physbeh.2017.03.040

Holzer, P. (1991). Capsaicin: cellular targets, mechanisms of action, and selectivity for thin sensory neurons. Pharmacol Rev, 43(2), 143–201.

Kakade, S., & Dayan, P. (2002). Dopamine: Generalization and bonuses. Neural Networks, 15(4-6), 549–559. https://doi.org/10.1016/S0893-6080(02)00048-5

King, C. D., Goodin, B., Kindler, L. L., Caudle, R. M., Edwards, R. R., Gravenstein, N., … Fillingim, R. B. (2013). Reduction of conditioned pain modulation in humans by naltrexone: An exploratory study of the effects of pain catastrophizing. Journal of Behavioral Medicine, 36(3), 315–327. https://doi.org/10.1007/s10865-012-9424-2

Kleinböhl, D., Hölzl, R., Möltner, A., Rommel, C., Weber, C., & Osswald, P. M. (1999). Psychophysical measures of sensitization to tonic heat discriminate chronic pain patients. Pain, 81(1-2), 35–43. https://doi.org/10.1016/S0304-3959(98)00266-8

Kroemer, N. B., Lee, Y., Pooseh, S., Eppinger, B., Goschke, T., & Smolka, M. N. (2019). L-DOPA reduces model-free control of behavior by attenuating the transfer of value to action. NeuroImage, 186, 113–125. https://doi.org/https://doi.org/10.1016/j.neuroimage.2018.10.075

Krohne, H. W., Egloff, B., Kohlmann, C. W., & Tausch, A. (1996). Untersuchungen mit einer Deutschen version der “Positive and Negative Affect Schedule” (PANAS). Diagnostica, 42(2), 139–156.

Kruschke, J. K. (2014). Doing Bayesian data analysis: A tutorial with R, JAGS, and Stan, second edition. https://doi.org/10.1016/B978-0-12-405888-0.09999-2

Kuznetsova, A., Brockhoff, P. B., & Christensen, R. H. B. (2017). lmerTest Package: Tests in Linear Mixed Effects Models. Journal of Statistical Software, 82(13), 1–26. https://doi.org/10.18637/JSS.V082.I13

Lang, P. J. (1980). Self-assessment manikin. Gainesville, FL: The Center for Research in Psychophysiology, University of Florida.

Langdon, A. J., Sharpe, M. J., Schoenbaum, G., & Niv, Y. (2018). Model-based predictions for dopamine. Current Opinion in Neurobiology, 49, 1–7. https://doi.org/10.1016/j.conb.2017.10.006

Leknes, S., Brooks, J. C. W., Wiech, K., & Tracey, I. (2008). Pain relief as an opponent process: a psychophysical investigation. The European Journal of Neuroscience, 28(4), 794–801. https://doi.org/10.1111/j.1460-9568.2008.06380.x

Lenth, R. (2020). Emmeans: estimated marginal means. R Package Version 1.5.0. Retrieved from https://cran.r-project.org/package=emmeans

Lewandowski, D., Kurowicka, D., & Joe, H. (2009). Generating random correlation matrices based on vines and extended onion method. Journal of Multivariate Analysis, 100(9), 1989–2001. https://doi.org/10.1016/j.jmva.2009.04.008

Leyton, M., Boileau, I., Benkelfat, C., Diksic, M., Baker, G., & Dagher, A. (2002). Amphetamine-induced increases in extracellular dopamine, drug wanting, and novelty seeking: a PET/[11C]raclopride study in healthy men. Neuropsychopharmacology, 27(6), 1027–1035. https://doi.org/10.1016/S0893-133X(02)00366-4

Löffler, M., Levine, S. M., Usai, K., Desch, S., Kandić, M., Nees, F., & Flor, H. (n.d.). Corticostriatal circuits in the transition to chronic back pain: the predictive role of reward learning. Cell Reports Medicine.

Luce, R. D. (1959). Individual Choice Behavior. New York: Wiley.

Maruyama, C., Deyama, S., Nagano, Y., Ide, S., Kaneda, K., Yoshioka, M., & Minami, M. (2018). Suppressive effects of morphine injected into the ventral bed nucleus of the stria terminalis on the affective, but not sensory, component of pain in rats. European Journal of Neuroscience, 47(1), 40–47. https://doi.org/10.1111/ejn.13776

Matsumoto, M., & Hikosaka, O. (2009). Two types of dopamine neuron distinctly convey positive and negative motivational signals. Nature, 459(7248), 837–841. https://doi.org/10.1038/nature08028

Meier, I. M., Eikemo, M., & Leknes, S. (2021). The Role of Mu-Opioids for Reward and Threat Processing in Humans: Bridging the Gap from Preclinical to Clinical Opioid Drug Studies. Current Addiction Reports, 8(2), 306–318. https://doi.org/10.1007/s40429-021-00366-8

Navratilova, E., Xie, J. Y., Meske, D., Qu, C., Morimura, K., Okun, A., … Porreca, F. (2015). Endogenous opioid activity in the anterior cingulate cortex is required for relief of pain. Journal of Neuroscience, 35(18), 7264–7271. https://doi.org/10.1523/JNEUROSCI.3862-14.2015

Navratilova, E., Xie, J. Y., Okun, A., Qu, C., Eyde, N., Ci, S., … Porreca, F. (2012). Pain relief produces negative reinforcement through activation of mesolimbic reward-valuation circuitry. Proceedings of the National Academy of Sciences, 109(50), 20709–20713. https://doi.org/10.1073/pnas.1214605109

Nyholm, D., Lewander, T., Gomes-Trolin, C., Bäckström, T., Panagiotidis, G., Ehrnebo, M., … Aquilonius, S. M. (2012). Pharmacokinetics of levodopa/carbidopa microtablets versus levodopa/benserazide and levodopa/carbidopa in healthy volunteers. Clinical Neuropharmacology, 35(3), 111–117. https://doi.org/10.1097/WNF.0b013e31825645d1

Pedersen, M. L., Frank, M. J., & Biele, G. (2017). The drift diffusion model as the choice rule in reinforcement learning. Psychonomic Bulletin and Review, 24(4), 1234–1251. https://doi.org/10.3758/s13423-016-1199-y

Pessiglione, M., Seymour, B., Flandin, G., Dolan, R. J., & Frith, C. D. (2006). Dopamine-dependent prediction errors underpin reward-seeking behaviour in humans. Nature, 442(7106), 1042–1045. https://doi.org/10.1038/nature05051

Peters, J., & D’Esposito, M. (2020). The drift diffusion model as the choice rule in inter-temporal and risky choice: A case study in medial orbitofrontal cortex lesion patients and controls. PLoS Computational Biology, 16(4), 1–25. https://doi.org/10.1371/journal.pcbi.1007615

Pizzagalli, D. a., Evins, a. E., Schetter, E. C., Frank, M. J., Pajtas, P. E., Santesso, D. L., & Culhane, M. (2008). Single dose of a dopamine agonist impairs reinforcement learning in humans: Behavioral evidence from a laboratory-based measure of reward responsiveness. Psychopharmacology, 196(2), 221–232. https://doi.org/10.1007/s00213-007-0957-y

R Core Team. (2019). R: A Language and Environment for Statistical Computing. Vienna, Austria. Retrieved from https://www.r-project.org/

Ratcliff, R. (1978). A theory of memory retrieval. Psychological Review, 85(2), 59–108. https://doi.org/10.1037/0033-295X.85.2.59

Ratcliff, R., & Rouder, J. N. (1998). Modeling Response Times for Two-Choice Decisions. Psychological Science, 9(5), 347–356. https://doi.org/10.1111/1467-9280.00067

Raynor, K., Kong, H., Chen, Y., Yasuda, K., Yu, L., Bell, G. I., & Reisine, T. (1994). Pharmacological characterization of the cloned κ-, δ-, and p-opioid receptors. Molecular Pharmacology, 45(2), 330–334.

Rescorla, R. A., & Wagner, A. R. (1972). A theory of Pavlovian conditioning and the effectiveness of reinforcement and non-reinforcement. In A. H. Black & W. F. Prokasy (Eds.), Classical Conditioning. 2. Current Research and Theory. (pp. 64–69). New York: Appleton-Century-Crofts.

Rinne, U. K., Birket-Smith, E., Dupont, E., Hansen, E., Hyyppä, M., Marttila, R., … Presthus, J. (1975). Levodopa alone and in combination with a peripheral decarboxylase inhibitor benserazide (madopar®) in the treatment of Parkinson’s disease. Journal of Neurology, 211(1), 1–9. https://doi.org/10.1007/BF00312459

Roth, M., & Hammelstein, P. (2012). The Need Inventory of Sensation Seeking (NISS). European Journal of Psychological Assessment, 28(1), 11–18. https://doi.org/10.1027/1015-5759/a000085

Santesso, D. L., Evins, a. E., Frank, M. J., Schetter, E. C., Bogdan, R., & Pizzagalli, D. a. (2009). Single dose of a dopamine agonist impairs reinforcement learning in humans: Evidence from event-related potentials and computational modeling of striatal-cortical function. Human Brain Mapping, 30(7), 1963–1976. https://doi.org/10.1002/hbm.20642

Savage, S. W., Zald, D. H., Cowan, R. L., Volkow, N. D., Marks-Shulman, P. A., Kessler, R. M., … Dunn, J. P. (2014). Regulation of novelty seeking by midbrain dopamine D2/D3 signaling and ghrelin is altered in obesity. Obesity, 22(6), 1452–1457. https://doi.org/10.1002/oby.20690

Schultz, W. (2007). Multiple dopamine functions at different time courses. Annual Review of Neuroscience, 30, 259–288. Retrieved from http://www.ncbi.nlm.nih.gov/entrez/query.fcgi?cmd=Retrieve&db=PubMed&dopt=Citation&list_uids=17600522

Schultz, W. (2016). Dopamine reward prediction error coding. Dialogues in Clinical Neuroscience, 18(1), 23–32.

Seymour, B. (2019). Pain: A Precision Signal for Reinforcement Learning and Control. Neuron, 101(6), 1029–1041. https://doi.org/10.1016/j.neuron.2019.01.055

Sherdell, L., Waugh, C. E., & Gotlib, I. H. (2012). Anticipatory pleasure predicts motivation for reward in major depression. Journal of Abnormal Psychology, 121(1), 51–60. https://doi.org/10.1037/a0024945

Sirucek, L., Price, R. C., Gandhi, W., Hoeppli, M.-E., Fahey, E., Qu, A., … Schweinhardt, P. (2021). Endogenous opioids contribute to the feeling of pain relief in humans. Pain, 162(12), 2821–2831. https://doi.org/10.1097/j.pain.0000000000002285

Smith, K. S., Berridge, K. C., & Aldridge, J. W. (2011). Disentangling pleasure from incentive salience and learning signals in brain reward circuitry. Proceedings of the National Academy of Sciences of the United States of America, 108(27). https://doi.org/10.1073/pnas.1101920108

Stan Development Team. (2020). RStan: the R interface to Stan. Retrieved from http://mc-stan.org/

Sutton, R. S., & Barto, A. G. (1998). Reinforcement Learning: An Introduction. Cambridge, MA, USA: MIT Press.

Tindell, A. J., Berridge, K. C., Zhang, J., Peciña, S., & Aldridge, J. W. (2005). Ventral pallidal neurons code incentive motivation: Amplification by mesolimbic sensitization and amphetamine. European Journal of Neuroscience, 22(10), 2617–2634. https://doi.org/10.1111/j.1460-9568.2005.04411.x

Vaillancourt, D. E., Schonfeld, D., Kwak, Y., Bohnen, N. I., & Seidler, R. (2013). Dopamine overdose hypothesis: Evidence and clinical implications. Movement Disorders, 28(14), 1920–1929. https://doi.org/10.1002/mds.25687

Vehtari, A., Gabry, J., Magnusson, M., Yao, Y., Bürkner, P.-C., Paananen, T., & Gelman, A. (2020). loo: Efficient leave-one-out cross-validation and WAIC for Bayesian models. Retrieved from https://mc-stan.org/loo/

Vehtari, A., Gelman, A., & Gabry, J. (2017). Practical Bayesian model evaluation using leave-one-out cross-validation and WAIC. Statistics and Computing, 27(5), 1413–1432. https://doi.org/10.1007/s11222-016-9696-4

Vellani, V., De Vries, L. P., Gaule, A., & Sharot, T. (2020). A selective effect of dopamine on information-seeking. ELife, 9, 1–14. https://doi.org/10.7554/ELIFE.59152

Villemure, C., Slotnick, B. M., & Bushnell, M. C. (2003). Effects of odors on pain perception: deciphering the roles of emotion and attention. Pain, 106(1-2), 101–108.

Vo, A., Seergobin, K. N., Morrow, S. A., & MacDonald, P. A. (2016). Levodopa impairs probabilistic reversal learning in healthy young adults. Psychopharmacology, 233(14), 2753–2763. https://doi.org/10.1007/s00213-016-4322-x

Wall, M. E., Brine, D. R., & Perez-Reyes, M. (1981). The metabolism of naltrexone in man. NIDA Research Monograph, 28, 105–131. Retrieved from http://europepmc.org/abstract/MED/6790999

Watson, D., Clark, L. A., & Tellegen, A. (1988). Development and Validation of Brief Measures of Positive and Negative Affect: The PANAS Scales. Journal of Personality and Social Psychology, 54(6), 1063–1070. https://doi.org/10.1037/0022-3514.54.6.1063

Weerts, E. M., McCaul, M. E., Kuwabara, H., Yang, X., Xu, X., Dannals, R. F., … Wand, G. S. (2013). Influence of OPRM1 Asn40Asp variant (A118G) on [11C]carfentanil binding potential: preliminary findings in human subjects. International Journal of Neuropsychopharmacology, 16(1), 47–53. https://doi.org/10.1017/S146114571200017X

Wittmann, B. C., Daw, N. D., Seymour, B., & Dolan, R. J. (2008). Striatal Activity Underlies Novelty-Based Choice in Humans. Neuron, 58(6), 967–973. https://doi.org/10.1016/j.neuron.2008.04.027

World Medical Association. (2013). World Medical Association Declaration of Helsinki: Ethical Principles for Medical Research Involving Human Subjects. JAMA, 310(20), 2191–2194. https://doi.org/10.1001/jama.2013.281053

Xie, J. Y., Qu, C., Patwardhan, A., Ossipov, M. H., Navratilova, E., Becerra, L., … Porreca, F. (2014). Activation of mesocorticolimbic reward circuits for assessment of relief of ongoing pain: A potential biomarker of efficacy. Pain, 155(8), 1659–1666. https://doi.org/10.1016/j.pain.2014.05.018

Zald, D. H., Cowan, R. L., Riccardi, P., Baldwin, R. M., Ansari, M. S., Li, R., … Kessler, R. M. (2008). Midbrain dopamine receptor availability is inversely associated with novelty-seeking traits in humans. The Journal of Neuroscience: The Official Journal of the Society for Neuroscience, 28(53), 14372–14378. https://doi.org/10.1523/JNEUROSCI.2423-08.2008

